# Deciphering the role of brainstem glycinergic neurons during startle and prepulse inhibition

**DOI:** 10.1101/2023.05.04.538315

**Authors:** Wanyun Huang, Jose C. Cano, Karine Fénelon

**Affiliations:** Biology Department, University of Massachusetts Amherst, Life Science Laboratories, 240 Thatcher Road, Amherst, MA, 01002, U.S.A; Department of Biological Sciences, University of Texas at El Paso, 500 West University Avenue, El Paso, TX, 79912, U.S.A; Lead Contact; Current address: University of Rochester Medical Center Rochester, NY

**Keywords:** sensorimotor gating, prepulse inhibition, aging, electrophysiology, caudal pontine reticular nucleus, amygdala, glycine transporter 2, optogenetics, startle.

## Abstract

Prepulse inhibition (PPI) of the auditory startle response is the gold standard operational measure of sensorimotor gating. Affected by various neurological and neuropsychiatric illnesses, PPI also declines during aging. While PPI deficits are often associated with cognitive overload, attention impairments and motor dysfunctions, their reversal is routinely used in experimental systems for drug screening. Yet, the cellular and circuit-level mechanisms of PPI remain unclear, even under non-pathological conditions. Recent evidence shows that neurons located in the brainstem caudal pontine reticular nucleus (PnC) expressing the glycine transporter type 2 (GlyT2^+^) receive inputs from the central nucleus of the amygdala (CeA) and contribute to PPI via an uncharted pathway. Using tract-tracing and immunohistochemical analyses in GlyT2-eGFP mice, we reveal the neuroanatomical location of CeA glutamatergic neurons innervating GlyT2^+^ neurons. Our precise *in vitro* optogenetic manipulations coupled to field electrophysiological recordings demonstrate that CeA glutamatergic inputs do suppress auditory neurotransmission in PnC neurons but not via action on transmitter release from auditory afferents. Rather, our data is consistent with excitatory drive onto GlyT2^+^ neurons. Indeed, our PPI experiments *in vivo* demonstrate that optogenetic activation of GlyT2^+^ PnC neurons increases PPI and is sufficient to induce PPI, clarifying the crucial role of these neurons in young GlyT2-Cre mice. In contrast, in older GlyT2-Cre mice, PPI is reduced and not further altered by optogenetic inhibition of GlyT2^+^ neurons. We conclude that GlyT2^+^ PnC neurons innervated by CeA glutamatergic inputs are crucial for PPI and we highlight their reduced activity during the age-dependent decline in PPI.

**SIGNIFICANCE STATEMENT:** Sensorimotor gating is a pre-attentive mechanism that declines with age and that is affected by neuropsychiatric and neurological disorders. Prepulse inhibition (PPI) of startle commonly measures sensorimotor gating to assess cognitive and motor symptoms and to screen drug efficacy. Yet, the neuronal mechanisms underlying PPI are still unresolved, limiting therapeutic advances. Here, we identify brainstem glycinergic neurons essential for PPI using tract tracing, *in vitro* electrophysiology and precise *in vivo* optogenetic manipulations during startle measurements in mice. Innervated by amygdala glutamatergic inputs, we show that these glycinergic neurons are essential and sufficient to induce PPI in young mice. In contrast, these neurons do not contribute to PPI in older mice. We provide new insights to the theoretical construct of PPI.

## INTRODUCTION

In mammals, startle responses are elicited by the sudden and strong stimulation of auditory or trigeminal inputs, leading to the contraction of skeletal and facial muscles (Yeomans et al. 2002; Yeomans and Frankland 1995). The acoustic startle reflex (ASR) can be inhibited by the presentation of a non-startling stimulus leading to prepulse inhibition (Graham, 1975; Hoffman and Ison, 1980; Koch, 1999).

PPI is the gold standard operational measure of sensorimotor gating, a fundamental pre-attentive mechanism by which sensory events can inhibit motor outputs. PPI is a translational assay used in humans (Kohl et al., 2013; Takahashi and Kamio, 2018) and other vertebrates (Koch, 1999; Burgess and Granato, 2007; Curtin and Preus, 2015; Aguilar et al., 2018). A hallmark of schizophrenia (Braff et al., 1978; Mena et al., 2016; Swerdlow and Geyer, 1998), PPI deficits are seen in other psychiatric and neurological disorders including autism spectrum disorders and obsessive compulsive disorder (Swerdlow et al., 1995; Castellanos et al., 1996; Braff et al., 2001). Additionally, in humans and rodents PPI declines with age by an unknown underlying mechanism (Swerdlow et al., 2023). PPI deficits are associated with cognitive symptoms including attention impairments and behavioral dysfunctions (Swerdlow et al., 1992). Currently, therapeutic advances rely on drug screening using the reversal of pharmacologically-induced PPI deficits in rodents (Frau et al., 2014; Lally et al., 2016a, b). Yet, common pharmaceuticals show inconsistent effects on PPI in affected individuals. This is because our fundamental understanding of the neural mechanisms regulating startle inhibition is still incomplete.

The mammalian primary startle pathway consists of spiral ganglion cells that innervate cochlear root neurons which transmit auditory information predominantly to contralateral giant neurons located in the brainstem caudal pontine reticular nucleus (PnC; **Fig. 1**). Electrical stimulation and lesion experiments in rodents demonstrate that startle-mediating PnC giant neurons send monosynaptic glutamatergic inputs onto cervical and spinal motoneurons to produce a motor response (Davis et al., 1982). The heterogeneous cell population of the PnC also contains GlyT2^+^ neurons (Rampon et al., 1996; Zeilhofer et al., 2005). While glycine is a major inhibitory neurotransmitter in the brainstem, the contribution of PnC glycinergic neurotransmission during PPI remains controversial. On one hand, immunohistochemical assays, *in situ* hybridization studies and *in vitro* electrophysiological recordings confirmed that startle-mediating PnC giant neurons express glycine receptors (Rampon et al., 1996; Sato et al., 1991; Geis and Schmid, 2011; Curtin and Preuss, 2015). On the other hand, the activation of glycine receptors in the rat PnC *in vivo* failed to alter startle amplitude (Koch and Friauf, 1995).

**Figure 1.**
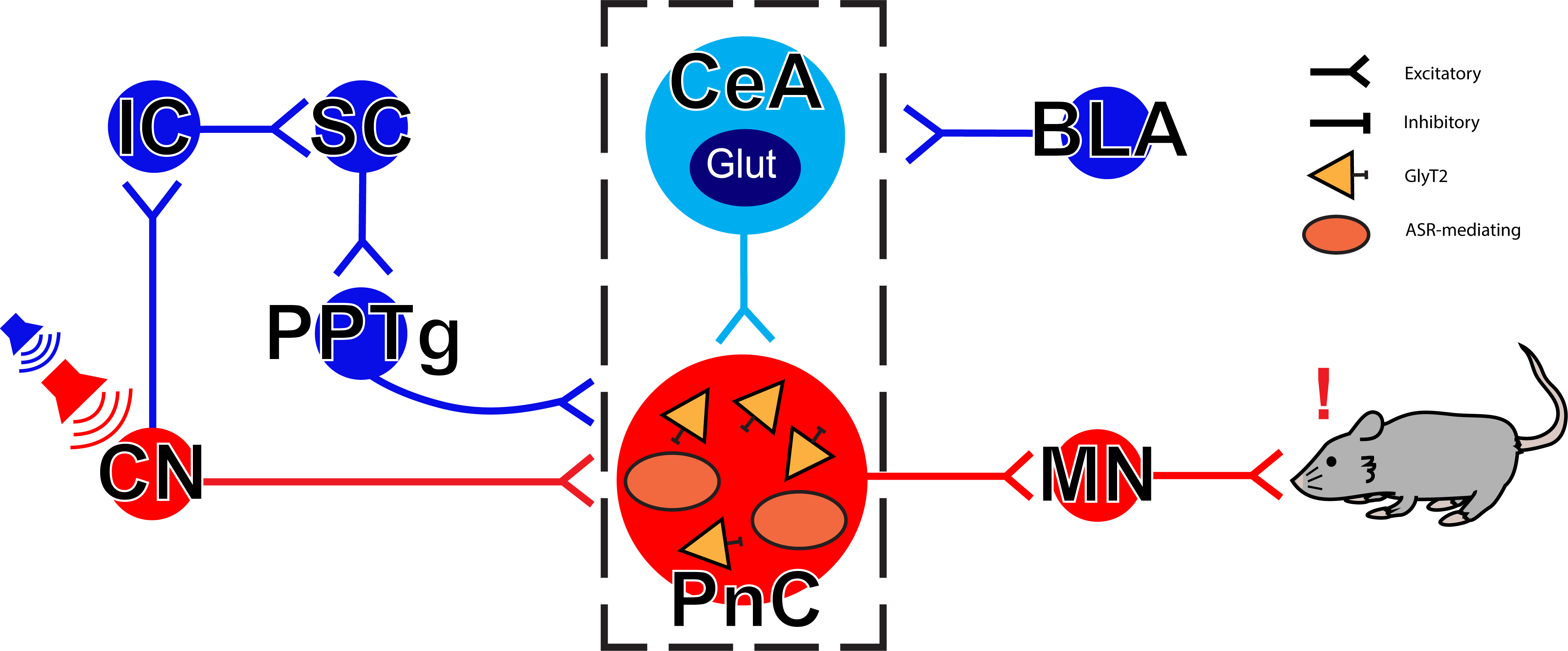
Simplified neuronal circuits underlying the acoustic startle response and PPI. The mammalian primary acoustic startle pathway (red pathway) includes primary auditory neurons that activate cochlear root and cochlear nuclei (CN) which then transmit auditory information mostly to contralateral startle-mediating giant neurons of the brainstem caudal pontine reticular nucleus (PnC). These PnC giant neurons directly activate cervical and spinal motor neurons (MNs), leading to a motor response. During PPI, the startle pathway has been thought to be inhibited by acoustic prepulses via an ascending pathway including the inferior (IC) and superior colliculi (SC), as well as the pedunculotegmental area (PPTg) (blue pathway). We recently showed that the central nucleus of the amygdala (CeA) contributes to PPI by sending glutamatergic inputs to PnC neurons expressing the glycine transporter type 2 (GlyT2^+^). BLA: basolateral amygdala; ASR: acoustic startle response. Modified from Cano et al., 2021.

Recent evidence ruled out the longstanding hypothesis that PPI is mediated by inhibitory cholinergic projections from the pedunculopontine tegmental nucleus (PPTg) to the PnC (MacLaren et al., 2014; Azzopardi et al., 2018; Fulcher et al., 2020). Using optogenetic inhibition, we recently showed in mice that aside from the corticostriatal-pallido-pontine pathway, GlyT2^+^ PnC neurons activated by glutamatergic monosynaptic inputs originating from the central nucleus of the amygdala (CeA), contribute to PPI (**Fig. 1;** Cano et al., 2021). But the neuroanatomy of the CeA-PnC connection and how it contributes to PPI is ill defined. Additionally, while the normal process of aging is associated with impaired inhibition (Hedden and Gabrieli, 2004; Perriol et al., 2005) and PPI decline (Terrasa et al., 2018), the contribution of GlyT2^+^ PnC neurons to PPI during aging is unknown.

Here, we used tract-tracing and immunohistochemical analyses to reveal the neuroanatomical distribution of CeA glutamatergic neurons and axon fibers innervating GlyT2^+^ neurons found ventro-medially in the PnC. We confirm that the CeA modulates auditory-evoked neurotransmission, using *in vitro* field electrophysiological recordings in PnC neurons. Then, we clarify the importance of glycinergic inhibition by demonstrating that GlyT2^+^ PnC neurons are essential and sufficient to elicit PPI, by performing *in vivo* optogenetic activation of these neurons in transgenic GlyT2-Cre mice during startle testing. Lastly, we highlight that the age-dependent decline in PPI is associated with a reduction in the activity of GlyT2^+^ PnC neurons.

## RESULTS

### CeA glutamatergic cells that innervate the PnC are predominantly found medially and laterally

Using retrograde and anterograde anatomical tracings in mice, we previously demonstrated that CeA glutamatergic neurons innervate the PnC and play an important role in startle modulation (Cano et al., 2021). While the CeA is typically considered a striatum-like GABAergic region, we showed that ~11% of CeA neurons project to the PnC and express both CamKIIα/eYFP and the vesicular glutamate transporter type 2 (VGluT2). However, the neuroanatomical distribution of these PnC-connected CeA glutamatergic neurons was not described, consistent with the fact that their function is largely overlooked. Here, to further understand how such a few number of glutamatergic neurons functionally impact the startle circuit, we aimed to determine the location of CeA neurons that innervate the PnC and whether these neurons are scattered or clustered within the CeA.

To map the anatomical distribution of CeA cells projecting into the PnC, we injected the retrograde tracer Fluorogold into the PnC of WT mice (**Fig. 2A, B**; C57BL/6 mice, N = 6). These injections targeted various sites, including the ventral, dorsal and lateral border regions of the PnC delineated by the 7^th^ cranial nerve. Mapped on the 73-78 levels of the Paxinos and Franklin Mouse Brain Atlas, we found that retrograde injection sites located in the ventro-medial portions of the PnC were the sites that back filled CeA cells most efficiently (**Fig. 2A,** filled circles). In contrast, dorsal or lateral PnC injections sites did not retrogradely label CeA neurons (**Fig. 2A;** open circles). No obvious cell cluster was identified in the CeA. Instead, Fluorogold-labeled cell bodies were found scattered throughout the medial and lateral regions of the CeA.

**Figure 2.**
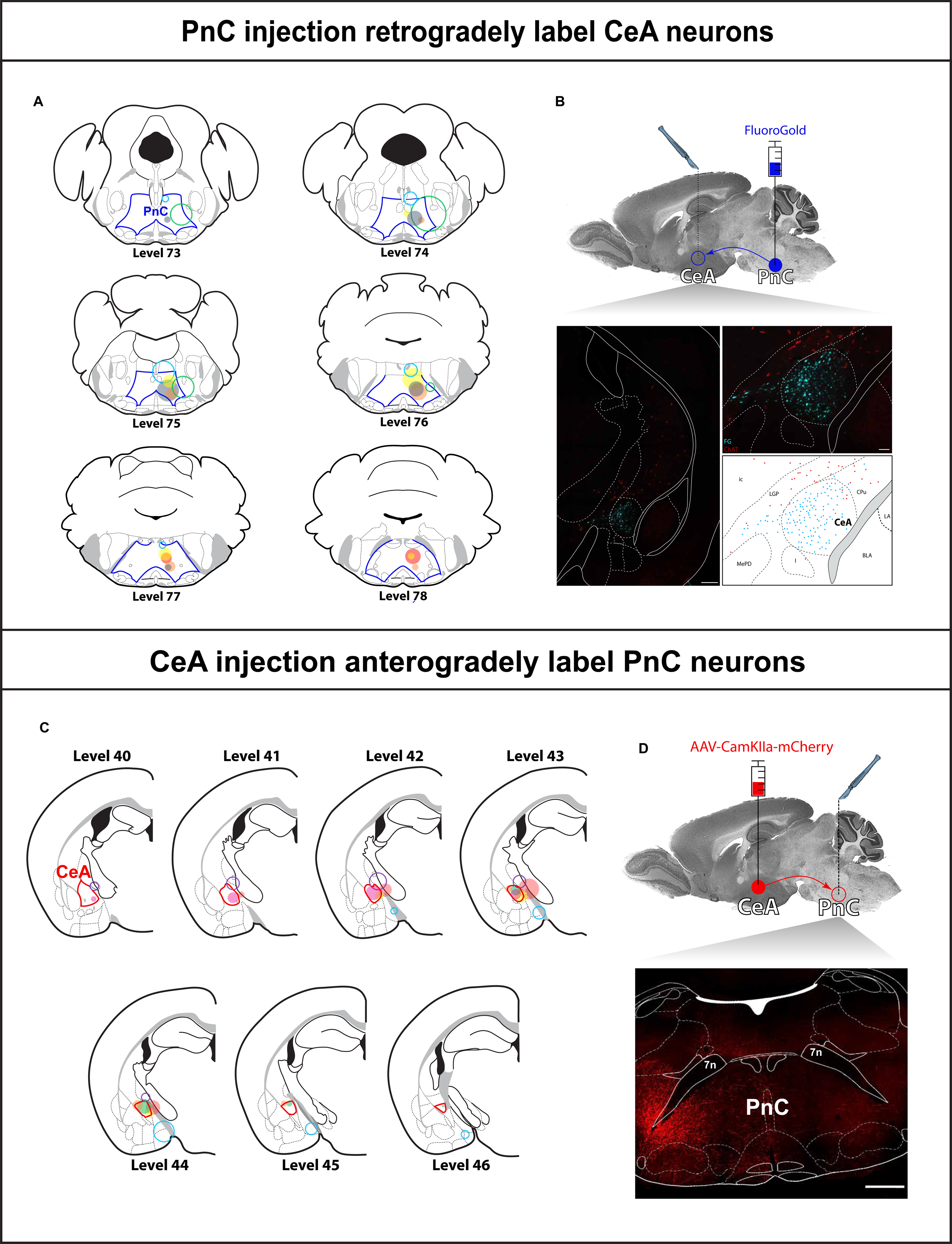
Distribution of PnC-connected CeA glutamatergic neurons. **(A)** Representative coronal PnC slices showing individual Fluoro-Gold injection sites that produced retrograde labelling (filled colored circles) or no retrograde labelling (open colored circles) in the CeA. The anteroposterior anatomical levels are from the Paxinos and Franklin Mouse Brain Atlas. PnC is delineated by solid blue lines. (**B**) *Top*, Mouse brain sagittal section illustrating the injection site of the retrograde tracer Fluoro-Gold in the PnC (filled blue circle) and the labeling site in the CeA (open blue circle) of WT mice. *Bottom left*, Representative image of a coronal brain section that includes retrogradely-labeled fluorescent Fluoro-Gold^+^ CeA neurons (cyan) and adjacent ChAT^+^ neurons (red) shown at low magnification. Fluoro-Gold labeled neurons were mainly observed within the CeA whereas ChAT^+^ neurons were found outside the CeA. Scale bar: 500μm. *Bottom right*, Representative image of Fluoro-Gold labeled CeA neurons and adjacent ChAT^+^ neurons mapped onto level 42 of the Paxinos and Franklin Mouse Brain Atlas and seen at higher magnification. Ic = insular cortex; LGP = lateral globus pallidus; MePD = posterodorsal medial amygdala; I = intercalated amygdaloid nuclei; CeA = Central nucleus of the amygdala; CPu = Striatum; LA = Lateral amygdala; BLA = basolateral amygdala. Scale bar: 100μm. N = 6 mice. (**C**) Representative coronal amygdala slices at different anteroposterior levels mapped on the Paxinos and Franklin Mouse Brain Atlas. Individual AAV injection sites in the CeA that resulted in mCherry-labeled axonal projections (filled colored circles) and AAV injection sites that resulted in no labeling (open colored circles) in the PnC are shown. CeA is delineated by solid red lines. (**D**) *Top*, Mouse brain sagittal section illustrating the injection site of an AAV-DJ-CamKIIα-mCherry virus in the CeA (filled red circle) and the projection site in the PnC (open red circle) of WT mice. *Bottom*, Representative image of a coronal PnC section delineated by the 7^th^ cranial nerves showing mCherry^+^ CeA axon labeling in the PnC, with a subset of CeA axons crossing the midline. N = 7 mice. Scale bar: 500μm.

Next, to visualize CeA glutamatergic axons in the PnC, we exclusively labeled PnC-projecting amygdala glutamatergic neurons by injecting the anterograde viral vector AAV-DJ-CamKIIα-mCherry, into the mouse amygdaloid complex (**Fig. 2C, D;** C57BL/6 mice, N = 7). Cells targeted by this intracranial viral injection were found within the CeA, as well as in areas medial and ventral to the amygdaloid complex. All cells were labeled in a section of 1-2 mm in the anterior-posterior axis, and approximately 30-50 cells were labeled at each injection site (**Fig. 2C**). To identify the location of CeA glutamatergic neurons projecting to the PnC, we examined the amygdala injection sites that produced anterograde mCherry labeling of CeA axons in the mouse PnC. Five out of the seven amygdala injection sites, mapped onto levels 40-46 of the Paxinos and Franklin Mouse Brain Atlas, produced anterograde labeling of CeA glutamatergic axons, in the PnC. These five injection sites included amygdala neurons confined within lateral and medial regions of the CeA (**Fig. 2C;** filled circles). Efficient CeA injection sites were found within the anterior-posterior axis of the CeA. The CamKIIα-mCherry^+^ CeA axonal projections that originated from these CeA injection sites, coursed in a lateral-to-medial fashion at the level of the PnC. In fact, these CeA projections coursed along the ventrolateral part of the principal sensory trigeminal nucleus (Pr5VL), the dorsomedial part of the principal sensory trigeminal nucleus (Pr5DM) and the alpha parvocellular reticular nucleus (PCRtA; lateral to the 7^th^ cranial nerve) before entering the PnC. While we previously reported that CeA glutamatergic axon fibers course ventrally within the PnC (dorsal to the superior olivary complex; Cano et al., 2021), here we also show that a subset of CeA glutamatergic axons clearly cross the PnC midline and some CeA glutamatergic fibers also ascend dorsally (**Fig. 2D; Additional file 1**).

The remaining two (out of seven) anterograde viral injections targeted cells located ventrally, outside the CeA and the amygdaloid complex (**Fig 2C;** open circles). As expected from previous rat studies (Rosen et al., 1991), these injections did not yield labeling in the mouse PnC. This indicates that the CamKII/mCherry^+^ fibers and terminals labeled in the PnC by viral injections into the CeA result from neuronal tracer uptaken by CeA glutamatergic neurons but not by other cells nearby the amygdaloid complex.

Overall, from these retrograde and anterograde labeling experiments, our results show for the first time that amygdala glutamatergic neurons that send inputs to the PnC are distributed within the medial and lateral regions of the CeA. Additionally, while their axons enter laterally in the PnC and then course within the ventro-medial portions of the PnC, we provide further evidence that some CeA glutamatergic axons cross the midline and can be found ascending dorsally.

### GlyT2^+^ neurons are densely distributed within the ventro-medial region of the PnC

To reconcile the fact that CeA neurons innervating the PnC startle circuit are excitatory but they contribute to an inhibitory phenomenon (i.e., PPI), we previously showed that CeA glutamatergic neurons innervate PnC neurons that express the glycine transporter type 2 (i.e., GlyT2^+^ PnC neurons; Cano et al., 2021). However, the anatomical organization of GlyT2^+^ neurons in the PnC was not provided. Therefore here, we wanted to show with more precision that GlyT2^+^ neurons are located ventro-medially in the PnC (Rampon et al., 1996; Zeilhofer et al., 2005; Giber et al., 2015), where numerous CeA glutamatergic fibers course and terminate.

To map the neuroanatomical distribution of GlyT2^+^ PnC neurons, we used levels 75-78 of the Paxinos and Franklin Mouse Brain Atlas, shown to contain the PnC (**Fig. 3**). Guided by the 7^th^ cranial nerve to delineate the PnC in GlyT2-eGFP mice (N = 4), we observed numerous GlyT2/eGFP^+^ neurons scattered throughout the extent of the PnC at the anatomical levels we assessed (**Fig. 3**). Several GlyT2/eGFP ^+^ neurons were clustered in close proximity to each other ventrally and along the midline of the PnC (**Fig. 3;** Level 75: 148 ± 6; Level 76: 228 ± 8; Level 77: 214 ± 11; Level 78: 194 ± 8 cell bodies). Additionally, as a second approach to visualize the distribution of GlyT2^+^ neurons in the PnC, we took advantage of mice that express the Cre recombinase enzyme under the control of the glycine transporter type 2 (i.e., GlyT2-Cre mice). To label GlyT2^+^ PnC neurons in GlyT2-Cre mice, we injected the Cre-dependent viral vector AAV-Ef1α-DIO-eYFP in the PnC (**Additional file 2**) and we observed a similar distribution of GlyT2^+^ PnC neurons as the results obtained from GlyT2-eGFP mice. Most importantly, we confirmed that GlyT2^+^ PnC neurons are densely located at sites innervated by CeA glutamatergic axonal projections, in the ventral part of the PnC. Overall, our labeling results reveal for the first time that GlyT2^+^ neurons located in the PnC are densely distributed along the midline and also dorsal to the olivary complex, medial to the 7^th^ cranial nerve.

**Figure 3.**
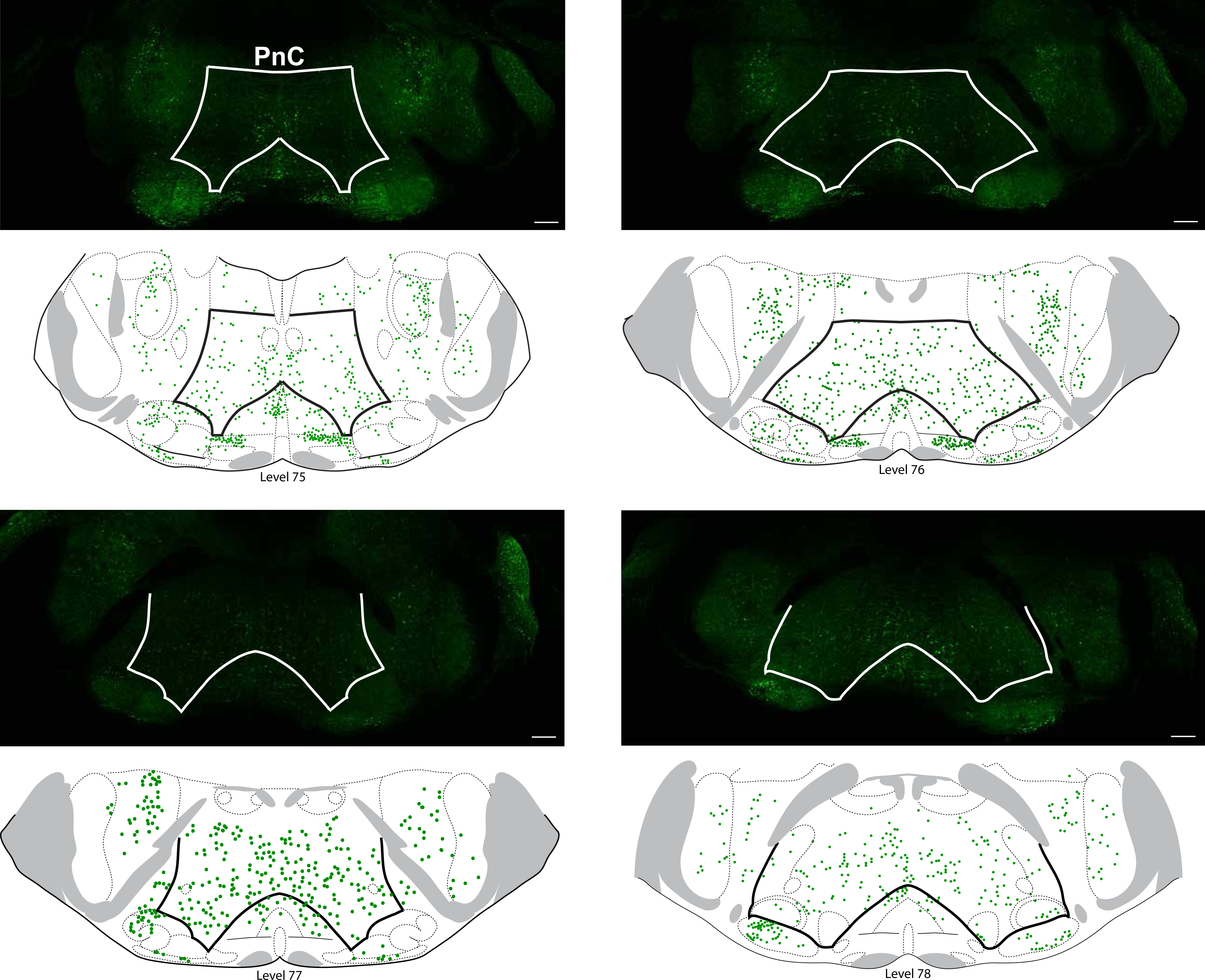
Distribution of GlyT2_+_ neurons in the PnC. *Top sections*, Representative coronal PnC sections showing the individual distribution of eGFP^+^ GlyT2 neurons (green) in an adult transgenic GlyT2-eGFP mouse, as visualized by immunofluorescence. PnC is delineated by solid white lines. *Bottom anatomical levels*, Distribution of GlyT2^+^ neurons (green) in the PnC mapped on anatomical level 75-78 of the Paxinos and Franklin Mouse Brain Atlas. PnC is delineated by solid black lines. Scale bars: 500µm. Representative of N = 6 mice.

### CeA glutamatergic neurons inhibit auditory neurotransmission in the PnC via a postsynaptic mechanism

Our neuroanatomical results described above confirm that in mice, CeA glutamatergic neurons innervate the PnC including GlyT2^+^ neurons. However, the synaptic mechanism by which CeA glutamatergic inputs modulate auditory neurotransmission in the PnC is unknown. Previous work in the zebrafish showed that during PPI, excitatory inputs inhibit auditory neurotransmission via a presynaptic mechanism within the acoustic startle circuit (Tabor et al., 2018). In fact, in the zebrafish, excitatory inputs act presynaptically on the terminals of auditory afferent fibers, and inhibit neurotransmitter release.

Therefore next, we aimed to identify the mechanism by which auditory neurotransmission is inhibited by CeA excitatory inputs in the PnC by performing *in vitro* extracellular field electrophysiological recordings in acute PnC slices from WT mice (**Fig. 4**). So first, we exclusively labeled CeA glutamatergic neurons and made them express Channelrhodopsin 2 (ChR2) by injecting the optogenetic viral vector AAV-DJ-CamKIIα-ChR2-eYFP into the CeA of WT mice. Following CeA unilateral intracranial injection, we obtained acute CeA and PnC slices. In mice injected with CamKIIα-ChR2-eYFP in the CeA, we ensured that CeA glutamatergic neurons successfully expressed ChR2. This was confirmed by recording field potentials in response to blue-light photo-stimulation of ChR2-eYFP^+^ CeA neurons (CeA slices) and ChR2-eYFP^+^ CeA axon fibers (PnC slices) (**Additional file 3**). Moreover, in PnC slices, blue-light evoked field excitatory post synaptic potentials (fEPSPs) were reversibly abolished by application of ionotropic glutamate receptor antagonists, AP5 (50 µM) and CNQX (25 µM) (**Additional file 4;** N = 7 mice, n = 21 slices), confirming the glutamatergic nature of the CeA-PnC connection, as we previously described using patch clamp recordings (Cano et al., 2021).

**Figure 4.**
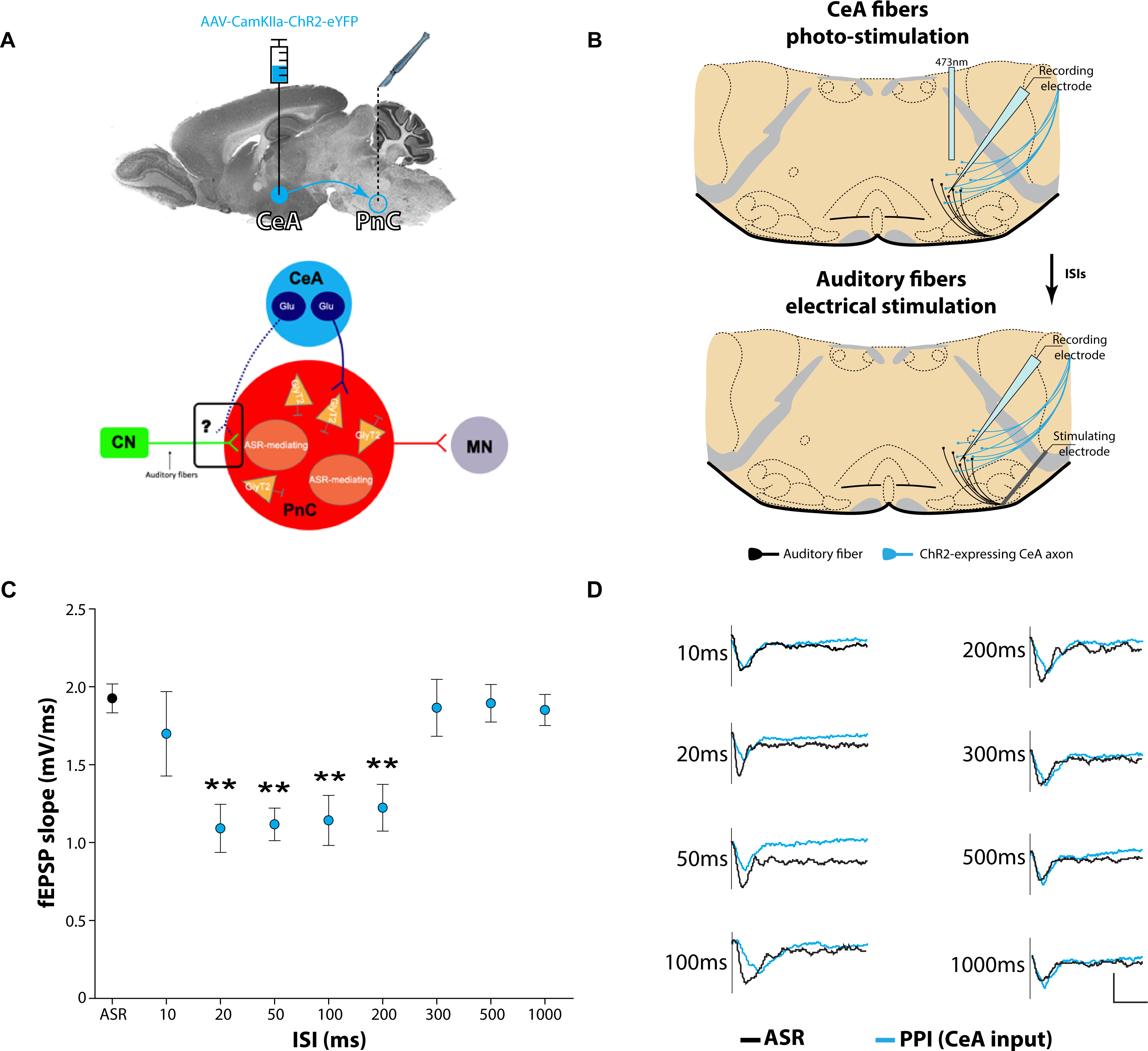
CeA glutamatergic inputs reduce auditory neurotransmission recorded *in vitro* in PnC neurons. **(A)** *Top*, Schematic illustration of a mouse brain sagittal section showing the injection of AAV-DJ-CamKIIα-ChR2-eYFP in the CeA (filled blue circle) of WT mice. PnC cut sections (open blue circle) were obtained following the AAV injection. *Bottom*, Schematic of the hypothesis being tested. **(B)** Schematic representation of a PnC slice used for *in vitro* field electrophysiological recordings. The blue optic fiber used to photo-activate ChR2-eYFP^+^ CeA glutamatergic axon fibers, the electrical stimulating electrode used to activate auditory fibers and the recording electrode used to record fEPSPs are illustrated. **(C)** Graph showing the mean slope of auditory-evoked fEPSPs recorded alone (“ASR”) or after the photo-activation of CeA glutamatergic inputs as a function of interstimulus intervals (ISIs) ranging from 10 to 1000ms. **(D)** Representative auditory-evoked fEPSPs recorded without (black traces) or recorded after the photo-activation of CeA glutamatergic inputs (blue traces) at various ISIs. N = 7 mice; n = 21 slices. Data are represented as mean ± SEM. \*\**p*<0.01.

To test whether photo-activating CeA glutamatergic inputs inhibits auditory neurotransmission in PnC slices, we electrically stimulated auditory fibers coursing dorsal to the lateroventral periolivary nucleus and recorded auditory neurotransmission as fEPSPs in PnC neurons (**Fig. 4A, B;** N = 7 mice, n = 21 slices). We hypothesized that if CeA glutamatergic neurons inhibit auditory neurotransmission, then optogenetic activation of CeA glutamatergic fibers would reduce auditory-evoked fEPSPs. As expected, our results show that photo-activating CeA fibers 20-200 milliseconds prior to the electrical stimulation of auditory fibers (**Fig. 4C, D**), significantly reduced auditory-evoked fEPSPs (F_(1,8)_ = 5.754, *p*<0.001). In contrast, the photo-activation of CeA fibers at very short or very long intervals prior to the stimulation of auditory fibers did not affect fEPSPs. These *in vitro* results are aligned with our previous *in vivo* findings showing that CeA-PnC glutamatergic synapses mediate PPI when the interval between prepulse and pulse stimuli ranges from 30-100 ms (Cano et al., 2021). Thus, photo-activation of CeA glutamatergic fibers coursing within the PnC startle circuit reduces auditory neurotransmission, consistent with their role during PPI *in vivo* at similar interstimulus intervals.

Then, to determine whether CeA glutamatergic inputs reduce auditory neurotransmission (i.e., fEPSPs) via a pre or post-synaptic mechanism in the PnC (relative to the auditory afferent fibers), we used the well-established paired pulse stimulation protocol. The paired pulse protocol uses two identical electrical stimuli applied at various inter-stimulus intervals (ISIs), to elicit two synaptic responses that can be compared. The resulting paired pulse ratio (PPR) has been shown to be inversely proportional to vesicle release probability following the first stimulus, allowing the detection of presynaptic effects (Xu-Friedman and Regehr, 2004). To distinguish between pre vs. post-synaptic effect, we recorded auditory-evoked paired fEPSPs in the absence and in the presence of the photo-activation of CeA glutamatergic fibers (**Additional file 5A;** N = 7 mice, n = 14 slices). Interestingly, our results show that PPR was not altered by the photo-activation of CeA glutamatergic fibers presented 50 milliseconds (**Additional file 5B, C;** 50ms: F_(1,7)_ = 0.510, *p* = 0.824) or 100 milliseconds (**Additional file 5D, E;** 100ms: F_(1,7)_ = 0.839, *p* = 0.559) prior to the paired fEPSPs elicited at various ISIs. Our findings indicate that CeA glutamatergic fibers do not act presynaptically on the synaptic terminals of auditory afferent fibers, in the PnC. Rather, our results suggest that CeA glutamatergic inputs reduce auditory neurotransmission in the PnC, via a post-synaptic mechanism.

### Optogenetic activation of PnC GlyT2_+_ neurons increases PPI

Our *in vitro* electrophysiological results above provide mechanistic insights further suggesting that during PPI, CeA glutamatergic inputs activate postsynaptic GlyT2^+^ PnC neurons which then inhibit auditory neurotransmission. We previously showed that inhibiting GlyT2^+^ PnC neurons *in vivo* decreases PPI (Cano et al., 2021) but the consequence of activating GlyT2^+^ PnC neurons during startle and PPI was not evaluated. Therefore next, we wanted to further determine the causal role of GlyT2^+^ PnC neurons *in vivo* during acoustic baseline startle and PPI.

To determine if photo-activating PnC GlyT2^+^ neurons concurrent with an acoustic startling pulse would alter baseline startle, we first made GlyT2^+^ neurons express ChR2. To do so, we injected the Cre-dependent excitatory optogenetic viral vector AAV-EF1α-double floxed-hChR2(H134R)-mCherry-WPRE-HGHpA in the Pnc of GlyT2-Cre mice. Following the PnC intracranial injection, an optic fiber was implanted ventro-medially in the PnC to photo-activate GlyT2^+^ neurons expressing ChR2, with blue light (**Fig. 5; N = 12 mice**).

**Figure 5.**
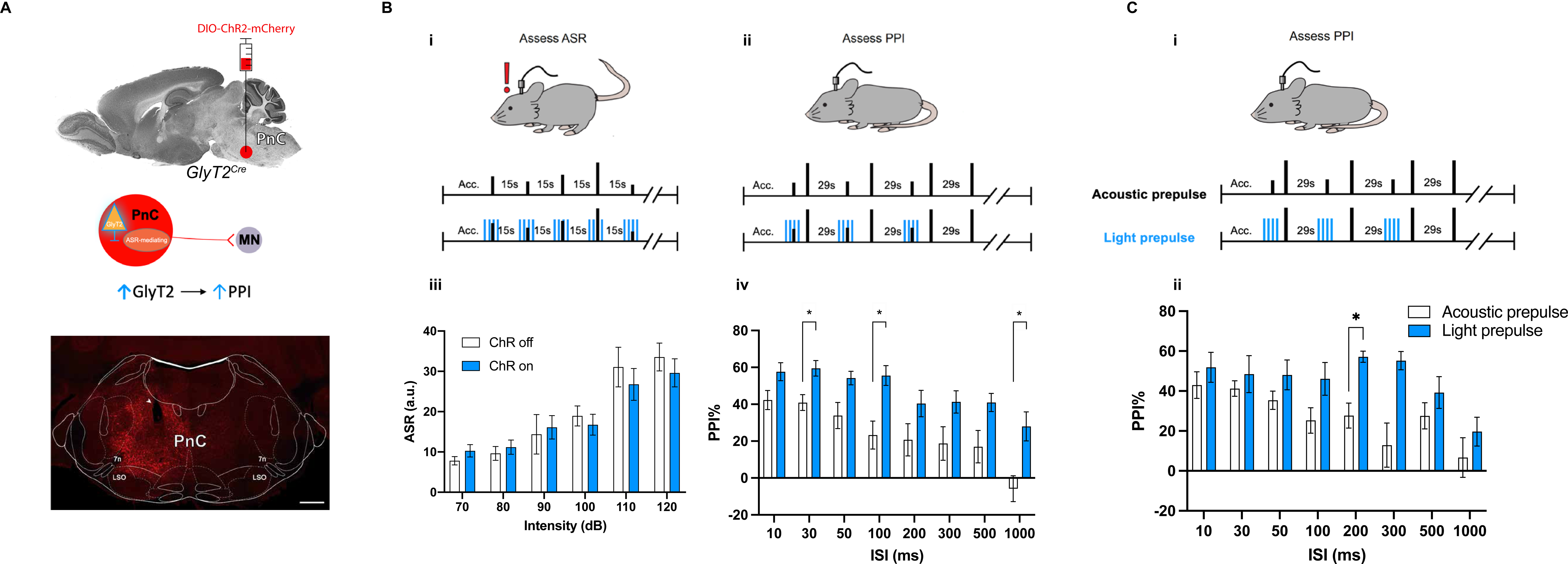
Photo-activation of GlyT2_+_ PnC neurons during ASR and PPI testing. **(A)** *Top*, Schematic illustration of the mouse brain sagittal section showing the injection site of the Cre-dependent and blue-light sensitive DIO-ChR2-mCherry virus in the PnC of GlyT2-Cre mice. *Middle*, the schematic of the hypothesis being tested. *Bottom*, Representative image showing the tract of implanted fiber optic (arrowhead) in the PnC. Scale bar, 500um; LSO, lateral superior olive; 7n, 7^th^ cranial nerve. **(B) (i-ii)** Schematic of the acoustic startle reflex and PPI protocols performed in GlyT2-Cre mice injected with ChR2. **(iii)** Graph showing the mean startle response amplitude as a function of sound at increasing intensities, both in the absence (open white bars; “ChR off”) and in the presence (blue bars; “ChR on”) of blue light photo-activation of GlyT2^+^ PnC neurons. Baseline startle amplitude was not affected by the photo-activation of GlyT2^+^ PnC neurons during 70-120 dB pulse alone acoustic stimulations [light: F_(1,11)_ = 0.228, *p* = 0.643; sound intensity: F_(5,55)_ =23.009, *p* < 0.001; light x sound intensity interaction: F_(5,55)_ = 1.393, *p* = 0.269]. (**iv**) Graph showing mean PPI values as a function of interstimulus intervals between acoustic prepulse and pulse, both in the absence (open white bars; “ChR off”) and in the presence (blue bars; “ChR on”) of blue light photo-activation of GlyT2^+^ PnC neurons. Photo-activation of GlyT2^+^ PnC neurons paired with acoustic prepulses increased PPI at all tested ISIs between the prepulse and the pulse. We found a significant effect of light: F_(1,11)_ = 17.196, *p* = 0.002 and ISI: F_(7,77)_ = 13.625, *p* < 0.001 on PPI. Two-way RM ANOVA, N = 12. **(C) (i)**, Schematic of the PPI protocols performed using acoustic prepulses (*top*) or blue light prepulses (*bottom*) in GlyT2-Cre mice. **(ii)** Graph showing mean PPI values as a function of interstimulus intervals, by using either sound (open white bars; “Acoustic prepulse”) or blue-light photo-activation of GlyT2^+^ PnC neurons (blue bars; “Light prepulse”) as a prepulse. Photo-activation of GlyT2^+^ PnC neurons as prepulses elicited PPI values comparable to or higher than acoustic prepulses, at all ISIs tested [light: F_(1,10)_ = 11.3, *p* = 0.007; ISI: F_(7,70)_ =5.423, *p* = 0.002; light x ISI interaction: F_(7,70)_ = 1.709, *p* = 0.121]. Two-way RM ANOVA, N = 6. Data are represented as mean ± SEM. \**p* < 0.05.

To test if GlyT2^+^ neurons contribute to baseline startle, mice were exposed to acoustic stimulations of increasing sound levels, with or without the photo-activation of GlyT2^+^ neurons. GlyT2^+^ PnC neurons were photo-activated with short trains of blue light at 5Hz (3ms pulse every 200ms) delivered concurrently with a pulse-alone acoustic stimulation. In all mice tested, sound intensities louder than 85 dB elicited a measurable acoustic startle response, characterized by a whole-body flexor muscle contraction (Ouagazzal et al., 2006; Valsamis and Schmid, 2011). Importantly, photo-activation of GlyT2^+^ PnC neurons paired with auditory startling stimulation did not affect the amplitude of the startle reflex of GlyT2-Cre mice (**Fig. 5B iii**). These results confirm that activating GlyT2^+^ neurons during a startling stimulation does not alter baseline startle. Our observations are consistent with our previous *in vivo* results showing that photo-inhibiting GlyT2^+^ neurons does not alter the baseline acoustic startle response (Cano et al., 2021).

Then, to assess whether activating GlyT2^+^ PnC neurons alters PPI, we photo-activated GlyT2^+^ PnC neurons concurrently with a non-startling acoustic prepulse, which was presented at various inter-stimulus intervals (ISIs) prior to an acoustic startling pulse (**Fig. 5B ii)**. Photo-activating GlyT2^+^ PnC neurons during the acoustic prepulse increased PPI by 31.38%-128.52% at ISIs between 10ms-1000ms (Main effect of light: F_(1,11)_ = 17.196, *p* = 0.002) in GlyT2-Cre mice (**Fig. 5B iv**). Overall, our *in vivo* photo-activation results suggest that during PPI, increasing the activity of GlyT2^+^ PnC neurons prior to a startling stimulation can further inhibit a startle response.

We next wanted to further verify that activating GlyT2^+^ PnC neurons alone is sufficient to induce PPI. We hypothesized that if GlyT2^+^ PnC neurons are necessary and sufficient for PPI, then photo-activating them prior to a startling pulse-alone stimulation would mimic the effect of an acoustic prepulse and lead to PPI (**Fig. 5C i; N = 6**). Indeed, when GlyT2^+^ PnC neurons were photo-activated prior to a startling pulse-alone stimulation at intervals used for acoustic PPI, startle was efficiently reduced. That is, photo-activating GlyT2^+^ PnC neurons in the absence of an acoustic prepulse was sufficient to successfully elicit PPI across different time intervals (**Fig. 5C ii**). Interestingly, the level of light-induced PPI was greater than the sound-induced PPI using acoustic prepulses (Main effect of light: F_(1,10)_ = 11.3, *p* = 0.007) particularly at ISI of 200ms (t_(5)_ = 4.293, *p* = 0.029). Importantly, our results confirm that photo-activating GlyT2^+^ PnC neurons alone is sufficient to inhibit a subsequent startle response.

### The contribution of GlyT2_+_ neurons during PPI declines with aging

While age-related deficits in inhibitory neurotransmission have been widely documented, very few studies have provided mechanistic insights towards the understanding of PPI decline with age (Geyer et al., 2001; Young et al., 2010). Importantly, current evidence in humans and rodents highlights the influence of age-related alterations of the amygdala and inhibitory mechanisms within related startle regions in modulating PPI (Rohleder et al., 2016). However, the role of GlyT2^+^ PnC in age-related PPI decline is unknown. GlyT2^+^ neurons are located in the PnC of mice and humans (Giber et al., 2015) and are intermingled with startle-mediating PnC giant neurons (Koch and Friauf, 1995; Rampon et al., 1996; Zeilhofer et al., 2005), therefore we next wanted to determine whether a change in the activity of GlyT2^+^ neurons is a potential neural substrate underlying the reduction of PPI with aging.

To test the contribution of GlyT2^+^ PnC neurons to PPI during aging, we used young GlyT2-Cre mice (2-3 months old, N = 13) and compared them to older GlyT2-Cre mice (9-12 months old, N = 8). All mice were injected with the Cre-dependent inhibitory optogenetic viral vector AAV-DJ-EF1α-DIO-eArch3.0-eYFP in the PnC. This viral injection in GlyT2-Cre mice allowed us to specifically target GlyT2^+^ PnC neurons and make them express the green-light sensitive Archaerhodopsin-3 (Arch3.0). We also injected WT mice with the same viral construct in the PnC and used these mice as controls for comparisons. In all animals, we used a unilateral optic fiber chronically implanted in the PnC, to deliver light and photo-inhibit GlyT2^+^ neurons during baseline startle and PPI.

To confirm that PPI and startle reactivity decrease with age in our mice, as was previously reported in rodent and human studies (Young et al., 2010, Shoji et al., 2016), we first compared PPI values using a prepulse of 5 dB above background noise and a pulse of 120 dB pulse for both young and aged mice. As expected (**Fig. 6B**), aged GlyT2-Cre mice exhibited significantly reduced acoustic PPI in comparison to young GlyT2-Cre mice (Main effect of age: F_(1,19)_ = 12.08, *p* = 0.025). More specifically, acoustic PPI was significantly decreased at short ISIs between 10-50 milliseconds (10ms: t_(19)_ = 3.15, *p* = 0.042; 30ms: t_(19)_ = 3.17, *p* = 0.04; 50ms: t_(19)_ = 3.4, *p* = 0.024).

**Figure 6.**
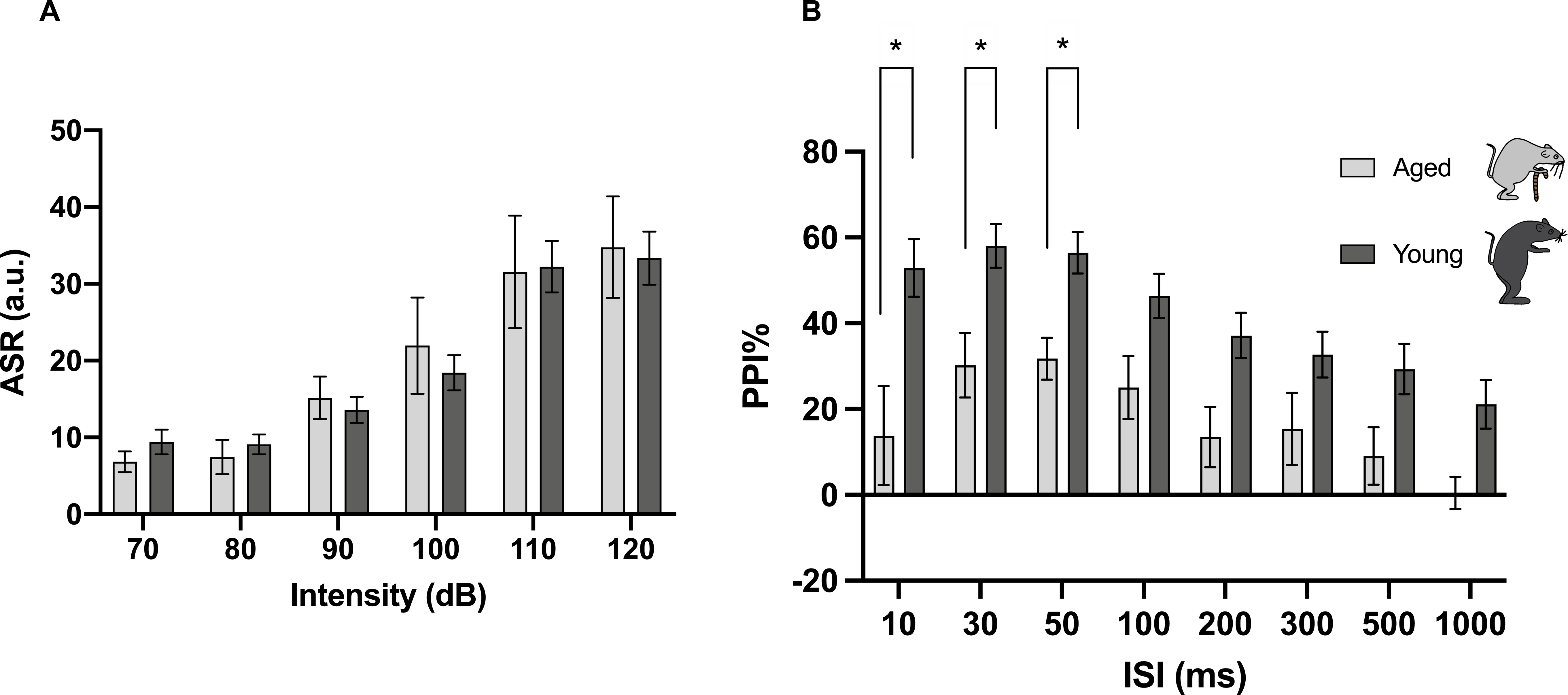
PPI levels decline with age. **(A)** Graph showing that mean startle amplitude as a function of sound intensity does not differ between young (2-3mo) and aged (9-12mo) GlyT2-Cre mice [age: F_(1,19)_ = 0.007, *p* = 0.932; sound intensity: F_(5,95)_ =37.817, *p* < 0.001; age x sound intensity interaction: F_(5,95)_ = 0.385, *p* = 0.664]. **(B)** Graph showing mean PPI levels significantly decrease in aged GlyT2-Cre mice at short ISIs between acoustic prepulse and pulse [age: F_(1,19)_ = 12.08, *p* = 0.025; ISI: F_(7,133)_ =14.18, *p* < 0.0001; age x ISI interaction: F_(7,133)_ = 1.148, *p* = 0.337]. Mixed ANOVA. Young GlyT2-Cre mice, N = 13 mice; Aged GlyT2-Cre mice, N = 8 mice. Data are represented as mean ± SEM. \**p* < 0.05. Images of mice icons are adapted from Ueta et al., 2020.

Then, to further determine whether the PPI reduction in the aged group was due to changes in baseline startle reactivity (**Fig. 6A**), we also compared startle reactivity between young and aged mice, at various sound levels (dB). Interestingly, there was no change in baseline startle reactivity among groups (Main effect of age: F_(1,19)_ = 0.007, *p* = 0.932). These data suggest that the reduction of PPI with aging is not correlated with baseline startle reactivity.

Next, to determine whether a reduction in the activity of GlyT2^+^ PnC neurons underlies the reduction in PPI in aged GlyT2-Cre mice, we used 3-ms pulses of green light stimulation delivered at 5 Hz, to photo-inhibit GlyT2^+^ PnC neurons during baseline startle in young and aged GlyT2-Cre mice. Similar to the results we previously reported in young GlyT2-Cre mice (Cano et al., 2021), the photo-inhibition of GlyT2^+^ PnC neurons concurrent with auditory stimulation of increasing intensity had no impact on startle responses in all mice tested (**Additional file 6**). Next, we used green light to photo-inhibit GlyT2^+^ PnC neurons during the prepulse and interstimulus interval of PPI trials. In young GlyT2-Cre mice, photo-inhibition of GlyT2^+^ PnC significantly decreased PPI by 37–40% at ISIs between 30–100 ms (30ms: t_(12)_ = 3.385, *p* = 0.043; 50ms: t_(12)_ = 3.779, *p* = 0.021; 100ms: t_(12)_ = 2.892, *p* = 0.031), as expected from our previous work. Interestingly, photo-inhibition of GlyT2^+^ PnC neurons had no effect on PPI in aged GlyT2-Cre mice (**Fig. 7**), similar to the results obtained from WT controls (**Additional file 7; N = 6 WT mice**). Overall, our results suggest that the PPI reduction observed in older GlyT2-Cre mice is associated with a reduction in the activity of GlyT2^+^ PnC neurons.

**Figure 7.**
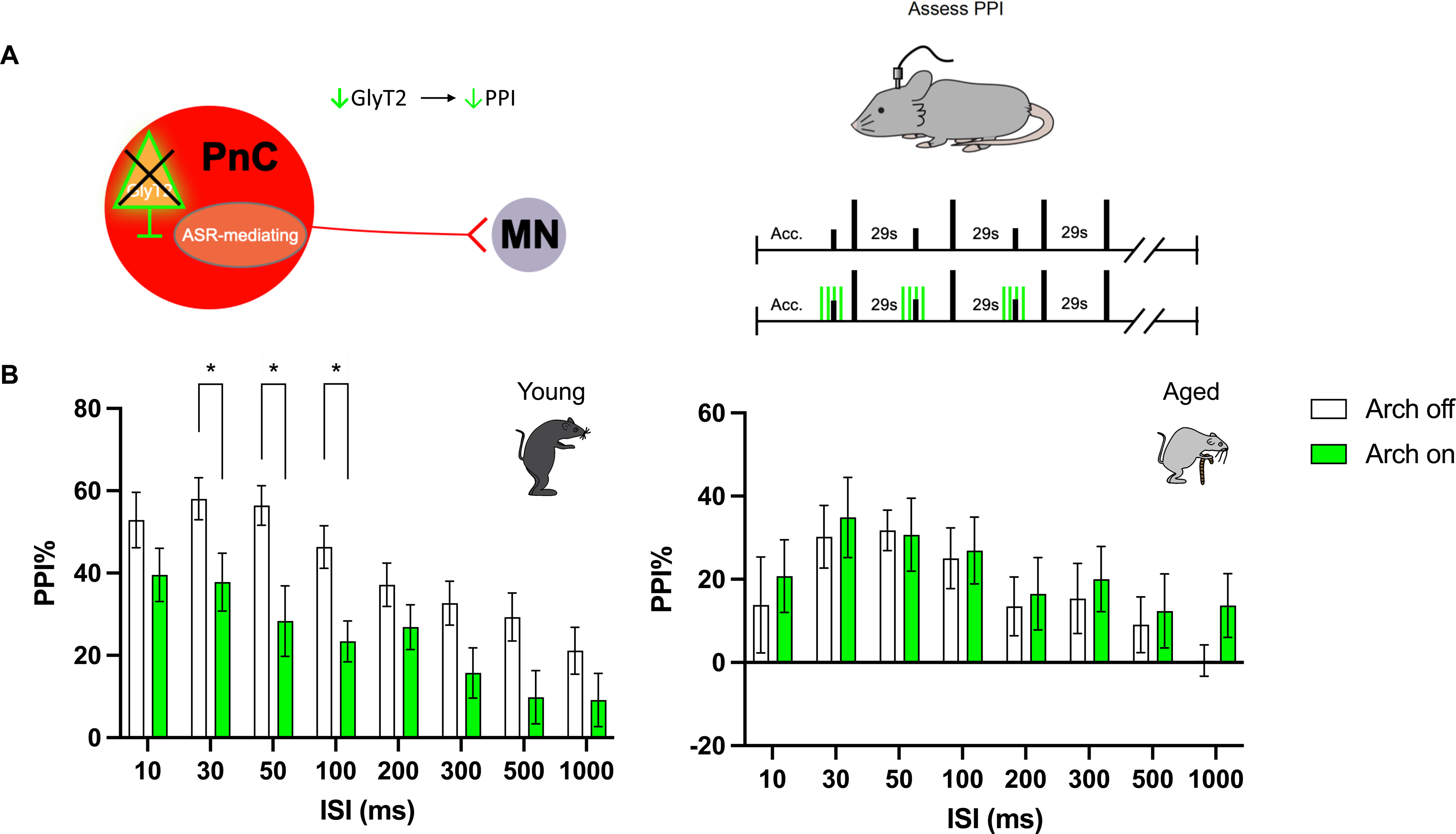
Photo-inhibition of GlyT2_+_ PnC neurons decreases PPI only in young mice. **(A)** *Left*, Schematic represents the hypothesis being tested. *Right*, Schematic of PPI protocols performed using young (2-3 mo) and aged (9-12 mo) GlyT2-Cre mice injected with the Cre-dependent and green light sensitive DIO-Arch-eYFP virus. **(B)** Graph showing mean PPI levels as a function of interstimulus intervals (ISI) between acoustic prepulse and pulse, both in the absence (open white bars; “Arch off”) and in the presence (green bars; “Arch on”) of green light photo-inhibition of GlyT2^+^PnC neurons. *Left*, Photo-inhibition of GlyT2^+^ PnC neurons paired with acoustic prepulses significantly decreased PPI at shorter ISIs in young mice [light: F_(1,24)_ = 10.14, *p* = 0.004]. *Right*, However, photo-inhibition of GlyT2^+^ PnC neurons had no effect on PPI in aged mice [light: F_(1,14)_ = 0.232, *p* = 0.637]. Two-way RM ANOVA. Young GlyT2-Cre mice, N = 13 mice; Aged GlyT2-Cre mice, N = 8 mice. Data are represented as mean ± SEM. \**p* < 0.05.

## DISCUSSION

Our results show that medial and lateral CeA glutamatergic neurons innervate GlyT2^+^ neurons located ventro-medially in the PnC. Additionally, our *in vitro* extracellular field electrophysiolgical results in PnC brain slices show that CeA glutamatergic inputs inhibit auditory neurotransmission but not by inhibiting neurotransmitter release from auditory afferents. We also show that photo-activating GlyT2^+^ PnC neurons increases PPI and their photo-activation alone is sufficient to induce PPI, further confirming their crucial role during PPI. Finally, we show that the contribution of GlyT2^+^ PnC neurons to PPI decreases with age. Overall, our results demonstrate that GlyT2^+^ PnC neurons innervated by CeA glutamatergic neurons are central to PPI, increasing our understanding of the mechanisms underlying PPI which declines during the normal process of aging.

### Neuroanatomy of CeA glutamatergic neurons innervating GlyT2_+_ PnC neurons

The CeA is considered a striatum-like structure, mainly consisting of GABAergic cells reminiscent of medium spiny neurons (McDonald and Augustine, 1993; Sun and Cassell, 1993; Pitkänen and Amaral, 1994; Saha et al., 2000). CeA local circuits have been shown to be composed of molecularly distinct populations with recurrent GABAergic inhibitory connections (Cassell et al., 1999; Ehrlich et al., 2009; Li et al., 2013; Kim et al., 2017; Wilson et al., 2019; Adke et al., 2021). Interestingly, tract tracing studies using VGAT-Cre mice confirmed that CeA GABAergic axons project to and innervate the brainstem (Liu et al., 2021), yet no GABAergic projection was found in the PnC. Despite the predominance of GABAergic neurons in the CeA, our results clearly indicate that CeA glutamatergic neurons are anatomically connected to the PnC as previously shown in rats (Rosen et al., 1991; Koch and Ebert, 1993; Lingenhöhl and Friauf, 1994), guinea pigs (Zhang et al., 2012) and supported by our recent findings in mice (Cano et al., 2021). Here, our retrograde and anterograde tracing experiments demonstrate that CeA glutamatergic fibers course ventro-medially within the PnC, where GlyT2^+^ neurons are located. We also show for the first time in mice that CeA glutamatergic axon fibers can cross the PnC midline and a subset of CeA fibers ascend to midbrain regions, as was recently suggested in rats and humans (Özkan et al., 2022). Since BLA fibers course through the CeA without projecting to the brainstem, our tracing results suggest that labelled fibers within the PnC resulting from amygdala injections reflect uptake from cells located in the CeA and not in the BLA. Finally, our injections outside the amygdaloid complex failed to produce labeling in the brainstem, as previously reported in rats (Rosen et al., 1991), further supporting a direct CeA-PnC glutamatergic connection.

### CeA glutamatergic inputs reduce auditory neurotransmission

To elucidate synaptic and cellular mechanisms underlying PPI in rodents, early studies focused on evaluating how brain regions innervating the PnC inhibit the activity of startle-mediating neurons. For example, *in vitro* electrophysiological experiments in rat brain slices demonstrated that the electrical stimulation of fibers originating from the pedunculopontine tegmental area (PPTg) reduces auditory neurotransmission recorded in PnC giant neurons (Homma et al., 2002; Bosch and Schmid, 2008). Here, we used highly specific optogenetics coupled to auditory fiber photostimulation to show that auditory-evoked fEPSPs in the PnC were reduced by photo-activation of CeA glutamatergic fibers presented 20-200ms earlier. This suggests that the CeA modulates auditory neurotransmission alongside other PnC inputs and/or neurotransmitter systems acting at other interstimulus intervals. In fact, evidence for inhibitory inputs additional to the glycinergic local neurons is provided by studies using *in vitro* bath application of GABA_A_, GABA_B_, ACh and glycine receptor agonists. These agonists reduce the amplitude of auditory-evoked EPSCs recorded in PnC giant neurons (Bosch and Schmid, 2006, 2008; Yeomans et al., 2010; Geis and Schmid, 2011), although the neuronal inputs driving release of GABA and ACh are not demonstrated.

One possible mechanism by which CeA neurons could inhibit auditory inputs into the PnC, separate from activation of the GlyT2+ neurons, is by depressing excitatory transmitter release from the auditory fibers themselves. Using paired pulse ratio, we tested for presynaptic modulation of auditory neurotransmission within the PnC by CeA glutamatergic fibers. Since paired-pulse ratios were not altered by the photo-stimulation of CeA glutamatergic fibers, our results suggest that the CeA inhibits auditory neurotransmission via a postsynaptic mechanism relative to auditory fibers. Additionally, previous *in vitro* rat studies showed that the amplitudes of the auditory-evoked EPSCs recorded in PnC giant neurons were not affected by the application of the glycine receptor antagonist strychnine, which excludes the possibility that auditory inputs within the primary startle pathway directly activate PnC glycinergic neurons leading to a feed forward inhibition of PnC giant neurons (Geis and Schmid, 2011). Together, these results are aligned with our hypothesis that a separate set of auditory inputs instead activate CeA glutamatergic neurons, which then activate postsynaptic PnC glycinergic neurons (Cano et al., 2021) leading to the inhibition of the PnC startle circuit. Such inhibitory modulation likely contributes to PPI at short interstimulus intervals alongside other neurotransmitter systems, as CeA glutamatergic fibers could also activate inhibitory mechanisms located in remote midbrain areas including the PPTg, shown to influence PPI at the PnC level via descending inputs (Bosch and Schmid 2008; Fulcher et al., 2020).

### Revisiting the contribution of GlyT2_+_ PnC neurons during PPI

We have recently showed in mice that CeA glutamatergic inputs activate GlyT2^+^ PnC neurons (Cano et al., 2021), and that these GlyT2^+^ PnC neurons, in turn, could inhibit startle-mediating PnC giant neurons known to express functional glycine receptors (Rampon et al., 1996; Sato et al., 1991; Zarbin et al., 1981; Geis and Schmid, 2011; Curtin and Preuss, 2015). While glycinergic neurons and axon fibers closely intermingle with PnC giant neurons (Rampon et al., 1996; Zeilhofer et al., 2005), few studies have investigated how glycine contributes to PPI. In early experiments, the initial assumption that glycine receptors underly PPI was not supported by an *in vivo* rat study that found no significant modification of startle responses or PPI after stereotactic injections of the glycine receptor agonist (β-alanine) or antagonist (Strychnine) into the PnC (Koch and Friauf, 1995). However, here we have revisited the function of glycine during PPI by directly activating GlyT2^+^ PnC neurons *in vivo* during PPI. Our results show that during PPI, photo-activating GlyT2^+^ PnC prior to a startling stimulation significantly attenuates sound-evoked startle responses and photo-activating these neurons alone at relevant interstimulus intervals induces PPI, demonstrating that GlyT2^+^ PnC neurons are sufficient to suppress a sound-induced startle response.

The contrasting findings between our work and that of others may reflect underlying differences in the experimental protocols used. In the Koch and Friauf (1995) study, the injected glycine receptor agonist might not have reached and activated glycine receptors expressed by startle-mediating PnC neurons at a sufficient concentration to inhibit them, so that no significant reduction of startle or PPI could occur (Koch and Friauf, 1995). Additionally, within the PnC, between 40-60 giant neurons among a total of 200-350 are reported to be immunopositive for glycine receptors (Koch et al., 1992). Yet, the proportion of PnC neurons (including non-giant neurons) responsive to glycine during PPI is still unclear. Interestingly, systemic administration of strychnine significantly increased startle, further demonstrating that glycine receptors contribute to startle modulation (Koch and Friauf, 1995). While we provided mechanistic evidence that PnC glycinergic neurons inhibit the PnC startle pathway, we cannot exclude the possibility that PnC glycinergic neurons also modulate upstream pathways including the intralaminar thalamic nuclei (Giber et al., 2015) shown to contribute to PPI (Yasuda et al., 2017) and behavioral arrest (Giber et al., 2015) relevant to aging and diseases (Kemether et al., 2003; Woodward et al., 2012).

### Effect of aging on baseline startle and PPI

From human studies (Blumenthal et al., 2006; Blumenthal and Swerdlow, 2002), it is known that aging is associated with a decline in ASR amplitude and with an inverted U-shaped function of PPI, where the lowest PPI values are seen in youngest and oldest subjects (Ellwanger et al., 2003). Additionally, current evidence suggests that age-related alterations of amygdala circuits influence PPI (Li et al., 2009; Rohleder et al., 2016; Reed et al., 2014; Sasse et al., 2014; Feng et al., 2011; Le Duc et al., 2016). While some strain differences exist (Ouagazzal et al., 2006), rodent studies support the effect of aging on both ASR and PPI measures seen in humans (Willott et al., 1994; Young et al., 2010). In wild-type C57BL/6 J mice, the ASR is highest at 2-3 months old and PPI values peak at 6–7 months old, after which they both gradually decrease with aging (Jafari et al., 2020; Ouagazzal et al., 2006; Shoji et al., 2016). Additionally, c-Fos expression data in rats recently highlighted the activation of the CeA during PPI (Tapias-Espinosa et al., 2019), confirming the involvement of amygdala-dependent inhibitory mechanisms during PPI (Cromwell et al., 2015; Webber et al., 2013). Here, we confirm for the first time that the activity of GlyT2^+^ PnC neurons, innervated by CeA glutamatergic inputs, decreases during aging. Further studies should address how GlyT2^+^ neurons, active during PPI, are affected by aging and further affected by neurological disorders.

## ACKNOWLEDGMENTS

We thank Ms. Andrea Gouin for the technical support, mice genotyping and for the UMass animal care; Dr. Stephanie Padilla (UMass) for the technical assistance; Dr. Joseph A. Gogos (Columbia University), Dr. Amy B. MacDermott (Columbia University), and Dr. Abigail Jensen (UMass) for the insightful discussions; and Dr. James Chambers for the assistance with the imaging performed at the Light Microscopy Facility at UMass Amherst.

## AUTHOR CONTRIBUTIONS

Conceptualization, W.H., J.C. and K.F.; Methodology, W.H., J.C. and K.F.; Investigation, W.H., J.C.; Formal Analysis, W.H., J.C. and K.F.; Writing - Original Draft, W.H., J.C. and K.F.; Writing - Review & Editing, W.H., J.C. and K.F.; Funding Acquisition, J.C. and K.F.; Resources, K.F.; Supervision, K.F.

## DECLARATION OF INTERESTS

The authors declare no competing interests.

**Additional file 1. Tracing of ascending CeA fibers.** Representative image of a coronal brain section containing the PPTg showing ascending CeA axons fibers labeled by red fluorescence. The red fluorescent tracer was injected unilaterally in the CeA of a WT mouse. The majority of ascending CeA fibers course ipsilaterally whereas a subset of CeA fibers cross the midline and ascend contralaterally to the injection side. PPTg is delineated by solid white lines. Scale bar: 500µm.

**Additional file 2. Distribution of GlyT2^+^ neurons in the PnC of GlyT2-Cre mice. (A)** Representative image of a coronal PnC section showing the distribution of eYFP^+^ GlyT2 neurons (green) in an adult transgenic GlyT2-Cre mouse. The Cre-dependent eYFP^+^ neuronal tracer was injected in the PnC, which is delineated by the 7^th^ cranial nerves. The density of GlyT2^+^ neurons is highest in the ventrol-medial PnC region. Scale bar: 500μm. **(B)** Higher magnification of GlyT2^+^ PnC neurons shown in A. Scale bar: 250μm.

**Additional file 3. The effect of blue light photo-stimulation on CeA glutamatergic neurons and axon fibers. (A)** *Top*, Schematic illustration of a mouse brain sagittal section showing the injection of AAV-DJ-CamKIIα-ChR2-eYFP in the CeA (filled blue circle) of WT animals. PnC cut sections (open blue circle) were obtained after AAV injection. *Middle*, Schematic representation of brain slices containing the CeA (left) or the PnC (right), respectively used for the *in vitro* optogenetic activation of CeA glutamatergic neurons or CeA glutamatergic axon fibers coupled to field electrophysiological recordings. The blue optic fiber used to photo-activate ChR2-eYFP^+^ CeA neurons or axon fibers and the recording electrode used to record fEPSPs are illustrated. *Bottom*, superimposed sample fEPSP traces elicited in response to blue-light photo-activation of ChR2-eYFP^+^ CeA neurons (left) or axon fibers (right) at increasing blue light intensities. **(B)** Graph showing the mean field potential slopes in response to the photo-activation of CeA glutamatergic neurons in CeA sections (blue circles) or CeA axon fibers in PnC sections (red circles) as a function of blue light intensity. The lack of responses recorded in brain sections from WT animals injected with the control virus AAV-DJ-CamKIIα-eYFP confirms opsin efficacy (gray open circles). N = 7 mice; n = 21 slices. Data presented as means ± SEM.

**Additional file 4. PnC neurons receive glutamatergic synaptic inputs from the CeA.** Graph showing the mean amplitude of *in vitro* fEPSPs elicited by the optogenetic activation of CeA glutamatergic axon fibers with blue light and recorded in PnC neurons in control, during the sequential bath application of AP5 and DNQX and following washout. These blue light-evoked fEPSPs recorded in the PnC were abolished by the bath-application of the glutamate receptor antagonist AP5 alone or AP5 + CNQX and recovered after washout. *Inset*: the schematic representation of a brain slice containing the PnC. The blue optic fiber used to photo-activate ChR2-eYFP^+^ CeA axon fibers, the recording electrode used to record fEPSPs in PnC neurons and the bath application of AP5 and CNXQ (green and yellow filled circles) are illustrated. *Bottom traces*: Superimposed light-evoked fEPSP traces (gray: all traces; black: averaged trace) recorded in the PnC under each experimental condition. Scale: 10 mV/15 ms. N = 7 mice, n = 21 slices. Data are represented as mean ± SEM. *P < 0.05, **P < 0.01.

**Additional file 5. CeA glutamatergic inputs do not affect the paired pulse ratio recorded in *in vitro* in PnC neurons. (A)** *Left*, Schematic illustration of the mouse brain sagittal section showing the injection site of AAV-CamKIIα-ChR2-eYFP in the CeA (filled blue circle) and the site of recording in the PnC (open blue circle). *Right*, In PnC coronal slices, photo-activation of ChR2-eYFP^+^ CeA glutamatergic axon fibers was delivered several milliseconds prior to two electrical auditory fiber stimulations separated by an interstimulus interval (ISI) of 50ms or 100ms. These electrical stimulations elicited two fEPSPs, and the ratio of fEPSP2/fEPSP1 was used to determine whether CeA glutamatergic fibers modulate PnC auditory neurotransmission via a pre or postsynaptic mechanism. *Inset*: presynaptic hypothesis being tested. **(B)** Graph showing the mean ratio of fEPSP2/fEPSP1 at an ISI of 50ms, both in the absence of (denoted as “Control”) or after the photo-activation of CeA glutamatergic fibers. **(C)** Representative paired fEPSPs recorded at ISI = 50 ms, in the absence or presence of CeA fibers photo-activation **(D)** Graph showing the mean ratio of fEPSP2/fEPSP1 at an ISI of 100ms, both in the absence of (denoted as “Control”) or after the photo-activation of CeA glutamatergic fibers. **(E)** Representative paired fEPSPs recorded at ISI = 100ms, in the absence or presence of CeA fibers photo-activation. The photo-activation of CeA glutamatergic fibers failed to modify fEPSP ratios, suggesting that CeA glutamatergic fibers modulate PnC auditory neurotransmission post-synaptically relative to auditory afferents. Scale in C, E: 0.2mV/ms. Representative of N = 7 mice, n = 14 slices. Data are represented as mean ± SEM. *P > 0.05, **P > 0.01.

**Additional file 6**. **Photo-inhibition of GlyT2_+_ PnC neurons does not affect baseline startle in both young and aged mice. (A)** *Left*, Schematic represents the hypothesis being tested. *Right*, Schematic of acoustic startle reflex protocols performed using young (2-3 mo) and aged (9-12 mo) GlyT2-Cre mice injected with the Cre-dependent and green light sensitive DIO-Arch-eYFP virus. (**A**) Graph showing the mean startle response amplitude as a function of sound at increasing intensities, both in the absence (open white bars; “Arch off”) and in the presence (green bars; “Arch on”) of green light photo-inhibition of GlyT2^+^ PnC neurons. Startle amplitude was not affected by the photo-inhibition of GlyT2^+^ PnC neurons during 70-120 dB acoustic pulses in both age groups. *Left*, startle responses in young adult mice [light: F_(1,24)_ = 0.244, *p* = 0.626; sound intensity: F_(5,120)_ = 61.49, *p* < 0.0001; light x sound intensity interaction: F_(5,120)_ = 0.411, *p* = 0.84]. *Right*, startle responses in aged adult mice [light: F_(1,14)_ < 0.001, *p* = 0.985; sound intensity: F(5,70) = 23.83, *p* < 0.0001; light x sound intensity interaction: F_(5,70)_ = 0.396, *p* = 0.85]. Two-way RM ANOVA. Young GlyT2-Cre mice, N = 13 mice; Aged GlyT2-Cre mice, N = 8 mice. Data are represented as mean ± SEM.

**Additional file 7. Green light does not affect baseline startle or PPI in WT control mice injected with the Cre-dependent Arch3.0 viral vector. (A)** *Left*, Schematic illustration of the mouse brain sagittal section showing the injection site of the Cre-dependent and green light sensitive DIO-Arch-eYFP virus in the PnC of WT mice used as controls. *Right*, Schematic of the acoustic startle reflex and PPI protocols performed in WT mice injected with Arch3.0. Rightmost is the schematic of the hypothesis being tested. **(B)** Graph showing the mean PPI as a function of interstimulus intervals, both in the absence (open white bars; “Arch off”) and in the presence (green bars; “Arch on”) of green light photo-inhibition of GlyT2^+^PnC neurons. Photo-inhibition of GlyT2^+^ PnC neurons did not occur in WT mice. Therefore, startle amplitude was not affected by green light presented during 70-120 dB pulse alone acoustic stimulations [light: F_(1,10)_ = 1.013, *p* = 0.338; sound intensity: F_(5,50)_ = 57.32, *p* < 0.0001; light x sound intensity interaction: F_(5,50)_ = 0.57, *p* = 0.723]. **(C)** Graph showing mean PPI values as a function of interstimulus intervals between acoustic prepulse and pulse, both in the absence (open white bars; “Arch off”) and in the presence (green bars; “Arch on”) of green light photo-inhibition. Green light did not inhibit GlyT2^+^ neurons in WT mice. Therefore, PPI was not affected by green light paired with acoustic prepulses at ISIs between 10 and 1000 ms [light: F_(1,10)_ = 0.103, *p* = 0.755; ISI: F_(7,70)_ = 7.548, *p* < 0.001; light x ISI interaction: F_(7,70)_ = 0.621, *p* = 0.737]. Two-way RM ANOVA. WT mice, N = 6. Data are represented as mean ± SEM.

## CONTACT FOR REAGENT AND RESOURCE SHARING

Further information and requests for reagents and other resources may be directed to and will be fulfilled by the Lead Contact, Karine Fénelon.

## MATERIALS and METHODS SUBJECT DETAILS

Experiments were performed on C57BL/6 male mice (N = 26; The Jackson Laboratory, Bar Harbor, ME), GlyT2-eGFP mice (N=6; graciously provided by Dr. Manuel Miranda-Arango, University of Texas at El Paso, El Paso, TX) and GlyT2-Cre^+/-^ mice (N = 33; graciously provided by Dr. Jack Feldman, University of California, Los Angeles). Litters were weaned at PND 21 and housed together until stereotaxic microinjections were performed. Mice received food and water *ad libitum* in a 12 hour light/dark cycle from 8:00 am to 8:00 pm. All the stereotaxic coordinates and cytoarchitectural boundaries were derived from Paxinos and Franklin Mouse Brain Atlas. Following surgical procedures, mice were single-housed and monitored for the duration of the recovery period. Experiments were performed in accordance with and approved by the Institutional Animal Care and Use Committee of the University of Texas at El Paso (UTEP) and the University of Massachusetts Amherst (UMass).

### Stereotaxic microinjections

Mice were sedated by inhaling 5% isoflurane vapors (Piramal Critical Care, Bethlehem, PA), then placed on a stereotaxic apparatus (model 900, David Kopf, Tujunga, CA) and immobilized using ear bars and a nose cone. Mice were maintained under 1.5%-2.5% isoflurane throughout the duration of the surgical procedure. With bregma as a reference, the head of the mice was leveled on all 3 axes. A craniotomy was performed directly dorsal to the injection site. Then, a microinjector (Stoelting Co., Wood Lane, IL) with a 5μl Hamilton syringe (Hamilton Company Inc., Reno, NV) and a 32 gauge steel needle were used to deliver viral vectors.

To target PnC (coordinates from vregma: AP -5.35mm; ML +0.5mm, DV +5.15mm) GlyT2^+^ neurons, 500nl of the AAV-Ef1α-double floxed-hChR2(H134R)-mCherry-WPRE-HGHpA (Addgene plasmid #20297; for optogenetic activation) viral vectors or AAV-Ef1α-DIO-eArch3.0-eYFP (Deisseroth Lab, virus #GVVC-AAV-055; for optogenetic inhibition) viral vectors were injected unilaterally in the PnC of C57BL/6 mice (N = 6) as controls or GlyT2-Cre mice (N = 33) expressing the Cre recombinase enzyme in GlyT2^+^ neurons. To label CeA neurons that project to the PnC, a separate WT cohort (N = 6) were unilaterally injected with 50-80nl of the retrograde neuronal tracer Fluoro-Gold (Molecular Probes, Eugene, OR, catalog #H22845, lot #1611168) in the PnC.

To target CeA (AP -1.35mm, ML +2.66mm, DV +4.6mm) glutamatergic neurons, 100-125nl of AAV-DJ-CamKIIα-mCherry (Deisseroth Lab, #GVVC-AAV-009; for anterograde tracing) viral vectors or AAV-DJ-CamKIIα-hChR2(H134R)-eYFP (Deisseroth Lab, #GVVCAAV-037, lot #3150; for electrophysiological recordings) viral vectors were injected unilaterally in the CeA of C57BL/6 mice (N = 14).

After viral injection, the microinjection syringe was left in place for 10 mins after infusion to limit spillover during needle retraction. Mice injected with Fluoro-Gold recovered for 5–7 days to allow optimal Fluoro-Gold retrograde transport to occur. AAV-injected mice recovered for 3–5 weeks to allow sufficient time for maximal viral transduction.

### Immunohistochemistry

Mice were perfused transcardially with 0.9% saline solution for 10 mins followed by 4% paraformaldehyde (PFA) in 0.1M phosphate buffer saline (PBS; pH 7.4) for 15 mins. The brains were then extracted, first post-fixed overnight in 4% PFA solution and next embedded in 30% sucrose in PBS for 2 days at 4 °C. The brains were then frozen in chilled hexanes for 1 min and stored at -80°C. Coronal sections (thickness = 30μm) were obtained using a microtome (Leica CM3050 cryostat) and then stored in cryoprotectant (50% 0.05 M phosphate buffer, 30% ethylene glycol, 20% glycerol) at -20°C.

For mice injected with Fluoro-Gold, coronal tissue sections containing the CeA were first washed with 0.1M Tris-buffered saline (TBS; pH 7.4), and incubated in blocking solution (2% normal donkey serum, 0.1% Triton X-100; in 0.1M TBS) for 1-2 hours at room temperature. Then tissue sections were incubated with a goat anti-ChAT primary antibody (1:100, Millipore, catalog #AB144P-200UL, lot #2854034) for 60 hours at 4°C, washed with TBS, and then incubated in a Cy3-conjugated donkey anti-goat secondary antibody (1:500, Jackson ImmunoResearch Laboratories, catalog #705-165-147, lot #115611) for 4 hours at room temperature. Similarly, for mice injected with AAV vectors containing eYFP, tissue sections at the level of PnC and the CeA were incubated in a chicken anti-GFP primary antibody (1:1000, Abcam, catalog #ab13970, lot #GR236651-13, RRID:AB_300798), then incubated in an Alexa Fluor 488-conjugated donkey anti-chicken secondary antibody (1:500, Jackson ImmunoResearch Laboratories, catalog #703-545-155, lot #130357). For mice injected with AAV vectors containing mCherry, tissue sections at the level of PnC and the CeA were incubated with a chicken anti-mCherry (1:1000, Abcam, catalog #ab205402, lot #GR225123-3, RRID:AB_2722769), then incubated with a Cy3-conjugated donkey anti-chicken secondary antibody (1:500, Jackson ImmunoResearch Laboratories, catalog #703-165-155, lot #130328, RRID: AB_2340363). After three rinses with TBS, brain slices were mounted and coverslipped to visualize injection and projection sites.

### Microscopy analysis

Tissue sections were analyzed by using an Axio Observer.Z1 epifluorescence microscope (Carl Zeiss Inc., Thornwood, NY) equipped with Fluoro-Gold, GFP, Cy3 filters, 10x, 20x, and 40x objectives, a motorized stage and Axiovision Rel. 4.8 software (Carl Zeiss Inc., Thornwood, NY). To create photomontages, single Z-plane images were obtained with the MosaiX module of the Axiovision Rel. 4.8 software at 10x or 20x for each fluorophore sequentially (1024 × 1024 pixel resolution). A total of 836 images (fluorescence and bright-field) were analyzed for each brain region and boundaries were delineated using Adobe Illustrator (Adobe, San Jose, CA).

### Electrophysiological recordings

Experiments were performed on adult mice 3-5 weeks after AAV-DJ-CamKIIα-hChR2(H134R)-eYFP injection in the CeA. The mice were anesthetized, followed by cervical dislocation and rapid decapitation. Then, the brain was harvested and placed in ice-cold dissecting solution with the following composition (in mM): sucrose (195), NaCl (10), KCl (2.5), NaH_2_PO_4_ (1), NaHCO_3_ (25), glucose (10), MgSO_4_ (4), CaCl_2_ (0.5), constantly bubbled with 5%CO_2_/95%O_2_. Then, 300μm-thick coronal sections containing the PnC or the CeA were cut using a vibratome (Leica VT 1200S, Wetzlar, Germany). These PnC or CeA sections were immediately transferred to a beaker filled with artificial cerebrospinal fluid (aCSF) at room temperature bubbled with 5%CO_2_/95%O_2_ with the following composition (in mM): NaCl (124), KCl (2.5), NaH_2_PO_4_ (1), NaHCO_3_ (25), Glucose (10), MgSO_4_ (1), CaCl_2_ (2). Slices were transferred to a field electrophysiological recording setup and allowed to recover on an interface chamber for at least 2 hours at 31–32°C. aCSF was fed by gravity at a rate of ≈2mL/min. The extracellular field electrophysiological signals were acquired using the pClamp10 software, the Digidata 1440A (Molecular Devices) and an extracellular amplifier (Cygnus Technologies). Using PnC or CeA slices ipsilateral to the viral injection site, field excitatory post-synaptic potentials (fEPSPs) were recorded via a glass microelectrode (3-5 MΩ; Warner Instruments, LLC, Hamden, CT) filled with aCSF, and placed in the lateroventral region of the PnC or the medial CeA. The fEPSPs were evoked with a blue LED (473 nm) delivered through a 200μm fiber optic mounted on a micromanipulator. The fiber optic was placed in close proximity to the recording electrode. The fEPSPs were quantified by measuring the initial slope of the peak negativity of the synaptic responses. Basal synaptic transmission was assessed within the CeA and at CeA-PnC synapses by gradually increasing the light intensity from 1-11 relative light units (1rlu = 0.43mW). The light pulse duration was 1ms. For the subsequent stimulation protocols, the stimulation intensity that generated a fEPSP equal to half of the maximum fEPSP obtained at maximum light intensity was used. Using that stimulation intensity, the probability of vesicle release at CeA-PnC excitatory synapses was assessed using paired light pulses delivered every 15 seconds with different interstimulus intervals (ISI in ms: 2, 5, 10, 20, 50, 100, 200, 300, 500, 1000 and 1500). Finally, to identify the neurotransmitter receptors involved at CeA-PnC excitatory synapses, the glutamatergic receptor antagonists, AP5 (50μM) and CNQX (25μM), were bath-applied, (i.e., added to the aCSF solution) for 20 mins before fEPSPs recordings. The drugs were stored as stock solutions in de-ionized water at -20 ℃, then freshly diluted in aCSF at the mentioned concentrations prior to use.

Auditory fibers were electrically stimulated with a bipolar stimulating electrode (tip diameter 0.2 mm, FHC, Bowdoin, ME, catalog# CBBRC75) placed on the LVPO (ventral to the LSO, ∼1.25mm ML) of acute slices at the level of the PnC (Bosch and Schmid, 2008). Similar to the fEPSP recorded in response to the photo-stimulation above, the recording electrode was placed in the lateroventral area of the PnC. Auditory-evoked fEPSPs were recorded in PnC sections. Basic synaptic transmission of auditory fibers in the PnC was characterized at 0.033 Hz, with increasing stimulation intensities and a pulse duration of 0.1ms. The subsequent protocols were performed at the stimulus intensity that elicited a auditory-evoked fEPSP one-third of the maximum fEPSP recorded.

Some experiments were performed with the recording electrode placed on the lateroventral PnC (just dorsal to the DPO and medial to the 7n), and with a blue optic fiber positioned to photo-activate CeA glutamatergic fibers coursing through the PnC. For these experiments, a 2ms light pulse was used to photo-activate CeA glutamatergic fibers, followed by a 0.1ms electrical stimulation that was used to electrically activate auditory fibers, in PnC cut sections. The light pulses preceeded the electrical stimulation by different ISI (in ms: 10, 20, 50, 100, 200, 300, 500, 1000, 1500) to determine at which ISI the CeA modifies auditory-evoked activity in the PnC. To determine whether CeA inputs modify auditory neurotransmission via a pre or post-synaptic mechanism, 1ms light pulse was delivered a few milliseconds (in ms: 5, 10, 20, 50, 100, 200, 300, 500, or 1000) before the paired electrical stimulations (separated by 50 or 100ms) of auditory fibers.

### Behavioral testing

Three weeks after the viral injection in the PnC, mice were sedated by inhaling 5% isoflurane vapors, placed in a stereotaxic apparatus, and immobilized using ear bars and a nose cone. Mice were maintained under anesthesia (1.5%-2% isoflurane), the head was leveled in all three axes. With bregma as a reference, a craniotomy was drilled directly dorsal to the implantation site. A cannula guide with a 200μm core optical fiber (Thorlabs, Newton, NJ) was then implanted at the level of the PnC (AP -5.35mm, ML +0.5mm, DV +4.75mm), and cemented to the skull with dental cement (Parkell, Edgewood, NY). Mice recovered for 7 days post-surgery before behavioral testing.

Behavioral testing of the acoustical startle response (ASR) and prepulse inhibition (PPI) were completed in an enclosed sound-attenuated startle chamber from PanLab (Harvard Apparatus, Holliston, MA). Sound pressure levels were calibrated using a standard SPL meter (model 407730, Extech, Nashua, NH). Mice were placed on a movement-sensitive platform. Vertical displacements of the platform induced by startle responses were converted into a voltage trace by a piezoelectric transducer located underneath the platform. Startle amplitude was measured as the peak to peak maximum startle magnitude of the signal measured during a 1s window following the presentation of the acoustic stimulation. Prior to any testing session, mice were first handled and acclimatized to the testing chamber, where the mice were presented to a 65dB background noise, for 10 mins. This acclimatization period was used to reduce the occurrence of movement and artifacts throughout testing trials. Following the acclimatization period, an input/output (I/O) assay was performed to test startle reactivity. This I/O test began with the presentation of a 40-ms sound at different intensities (in dB: 70, 80, 90, 100, 110, and 120) every 15s, in a pseudorandomized order. Background noise (65dB) was presented during the 15s between sounds. A total of 35 trials (i.e. 7 sound intensities, each sound presented 5 times) were acquired and quantified. Startle reactivity, derived from this I/O assay, allowed the gain of the movement-sensitive platform to be set. This gain allowed the startle responses to be detected within a measurable range. Once determined, the gain for each experimental subject was kept constant throughout the remaining of the experiment.

Following a one-hour resting period, mice were presented with seven startle-inducing 120 dB (40 ms) sounds called “Pulse-alone” stimulations. These 120 dB sounds were presented every 29s (interspersed with 65dB background noise) and were used to achieve a stable baseline startle response. The following PPI test consisted of two different conditions as follows: 1-startling pulse-alone stimulations (120 dB noise; 40 ms), and 2-combinations of a prepulse (75 dB noise; 20 ms) followed by a 120 dB startling pulse (40 ms) at 8 different interstimulus intervals (ISI, end of prepulse to the onset of startling pulse, in ms): 10, 30, 50, 100, 200, 300, 500, and 1000. The inter-trial interval of these two conditions was 29s. All the testing trials were designed and recorded using PACKWIN V2.0 software in PanLab System (Harvard Apparatus, Holliston, MA).

For combined optogenetic manipulations, mice injected with viral vectors containing either ChR2 (N=12 mice) or Arch3.0 (N=27 mice) were tested in the startle chamber. These animals were closely monitored to ensure that they were comfortably tethered to an optic fiber, which exited through a small opening from the roof of the startle chamber. The optic fiber (200 µm diameter, Thorlabs, Newton, NJ) was connected to a cannula implanted on the head of the mouse with a zirconia sleeve (Thorlabs, Newton, NJ). Animals were tethered ∼15 mins before testing and allowed to move freely, exploring their home cages before being transferred to the startle chamber. Optogenetic stimulation was triggered by a signal from the Packwin software which was transformed into a TTL-pulse. This TTL-pulse triggered a waveform generator (DG1022, Rigol Technologies) to modulate light stimulation. Photo-stimulation was delivered using a blue 473nm laser (Opto Engine LLC, Midvale, UT) for ChR2 activation, and photo-inhibition was delivered using a green 532nm LED (Plexon, Dallas, TX) for Arch3.0 activation.

During ASR trials paired with optogenetic manipulations, a train of light stimulation (3ms light ON, 197ms light OFF) was delivered at 5Hz 400ms before the pulse, lasting the entire pulse. During PPI trials paired with optogenetic manipulations, a train of light stimulation (3ms light ON, 197ms light OFF) was delivered at 5Hz 400ms before the prepulse, lasting the entire prepulse. During PPI trials with optical stimulation used as a prepulse, a 50Hz train of light stimulation (3 pulses of 15ms light ON, 5ms light OFF) was delivered at various ISI (10, 30, 50, 100, 200, 300, 500, and 1000 ms; end of prepulse to the onset of pulse) prior to the startling pulse. At the end of each experiment, histological analyses were performed to confirm that: 1-the injected viral particles were confined to the PnC, and 2-the cannula guide placement was successfully aimed at the PnC. If these criteria were not met, the subject was excluded from the study.

### Quantification and statistical analysis

Statistical analyses were performed using SigmaPlot (Systat Software, Inc., San Jose, CA). The assumptions of normality and equal variance of the data were first tested and an equivalent nonparametric test was run if assumptions were not met. For repeated-measures ANOVA, Mauchly’s test of sphericity was run first and either Greenhouse-Geisser or Huynh-Feldt corrections were applied if sphericity was violated. If posthoc tests were necessary, a Student’s t-test with Bonferroni correction was performed. The criterion for significance was 0.05.

For the results of extracellular field recordings with receptor antagonists, one-way repeated-measures ANOVA were used to assess the effect of the receptor antagonists on the light-evoked fEPSPs. For PPI *in vitro* results, one-way ANOVA were used to reveal if at any ISI the electrically-evoked fEPSPs were significantly attenuated by the optical stimulation of CeA-PnC excitatory synapses. For the results of the *in vivo* startle response experiments, two-way repeated-measures ANOVA was used assess the effect of sound intensity and optical manipulation on the I/O function of the startle response. For the results of the *in vivo* PPI experiments, two-way repeated-measures ANOVA was used to assess the effect of the ISI and and optical manipulation on PPI levels. Prepulse inhibition was expressed as a percentage of prepulse inhibition: %PPI = [1-(startle amplitude during “Prepulse+Pulse” trials/startle amplitude during “Pulse only” trials) × 100], which indicates the amount of startle suppression due to the presentation of prepulse. Mixed ANOVA was used to assess age effect on startle response/PPI between age groups. Sample sizes were chosen based on expected outcomes, variances, and power analysis. Data are presented as means ± SEM. N indicates the total number of animals.

