## Supplementary figures and images for "Deciphering the role of brainstem glycinergic neurons during startle and prepulse inhibition"

### Additional File 1

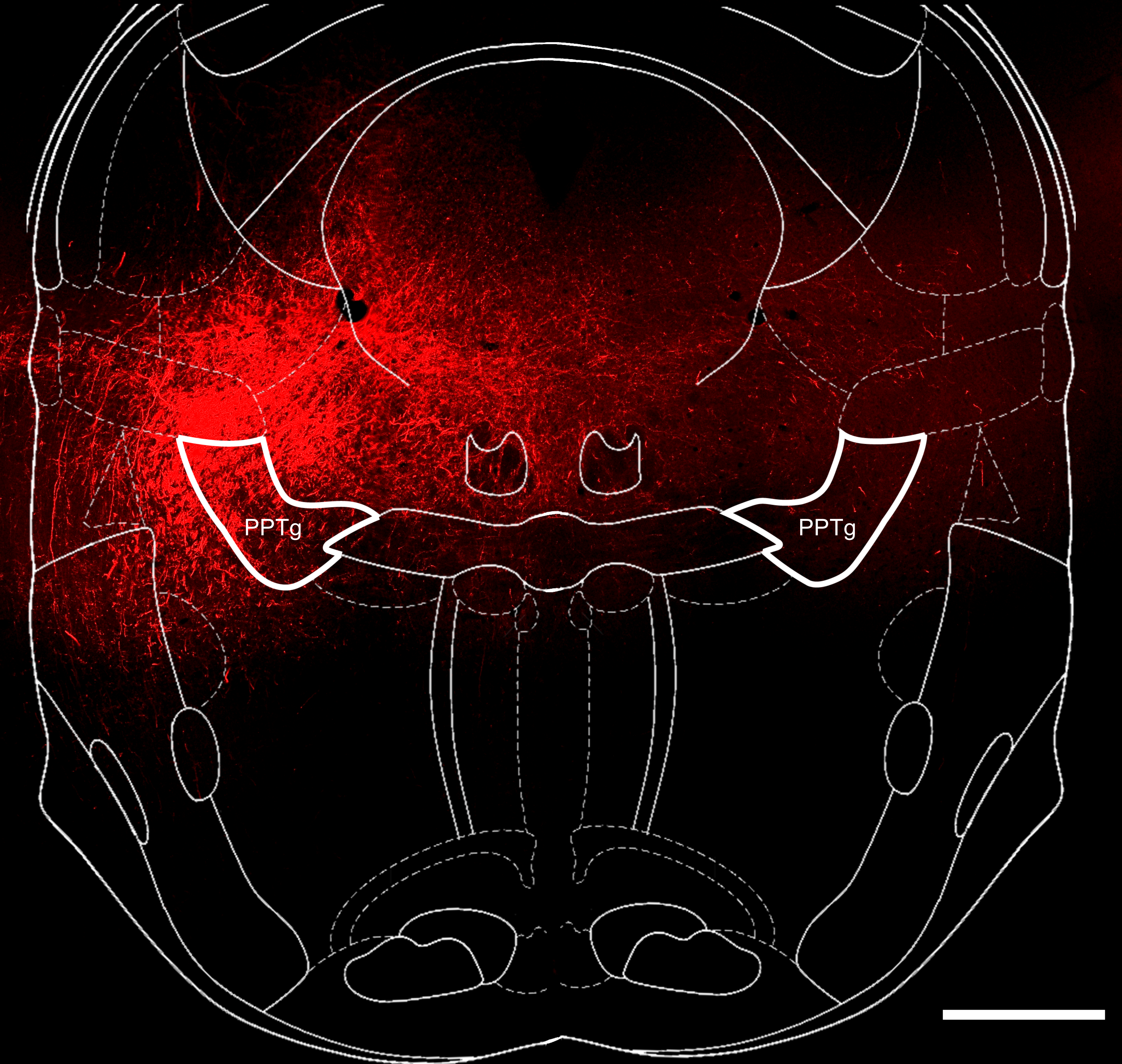

### Additional File 2

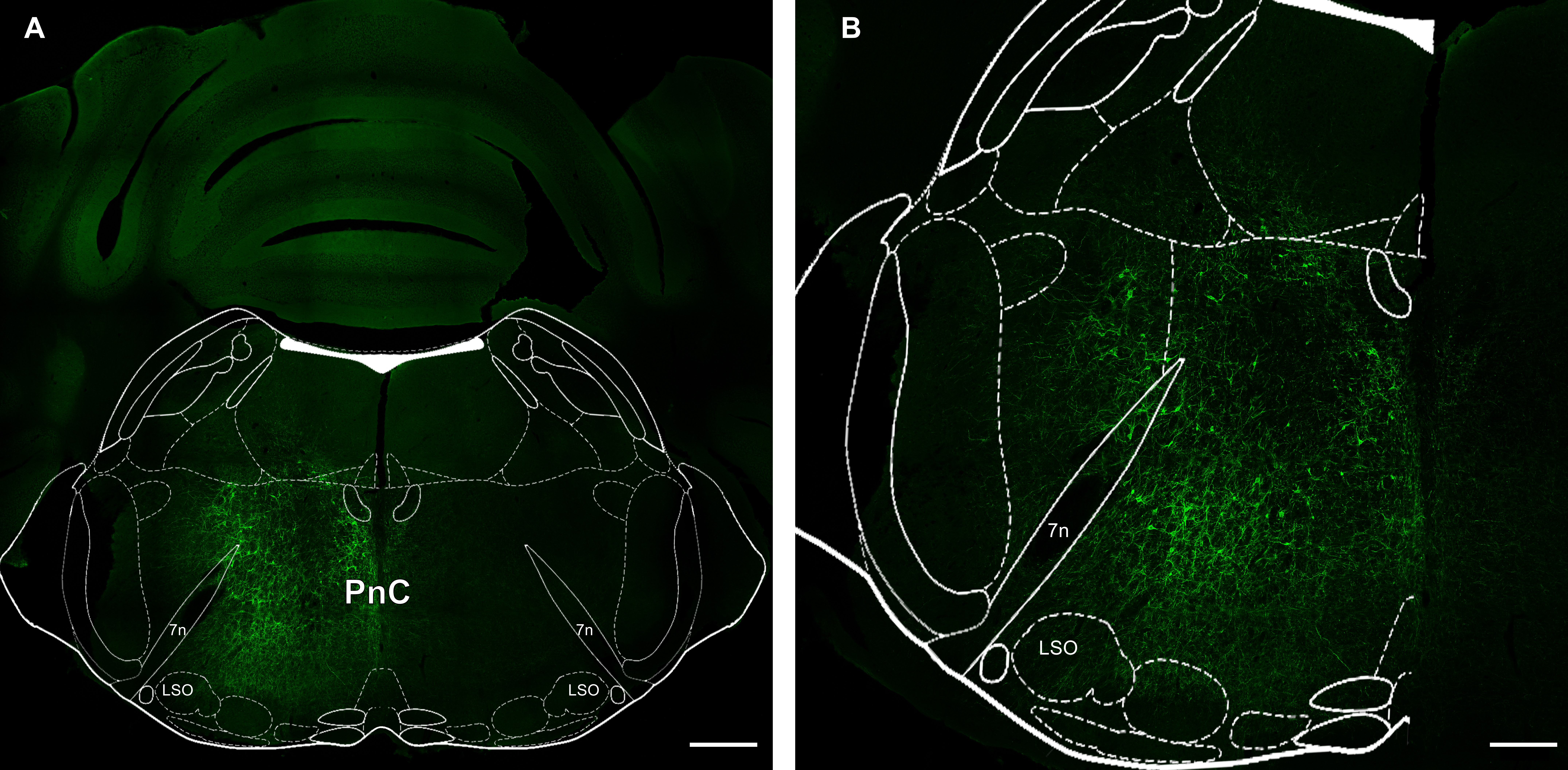

### Additional File 3

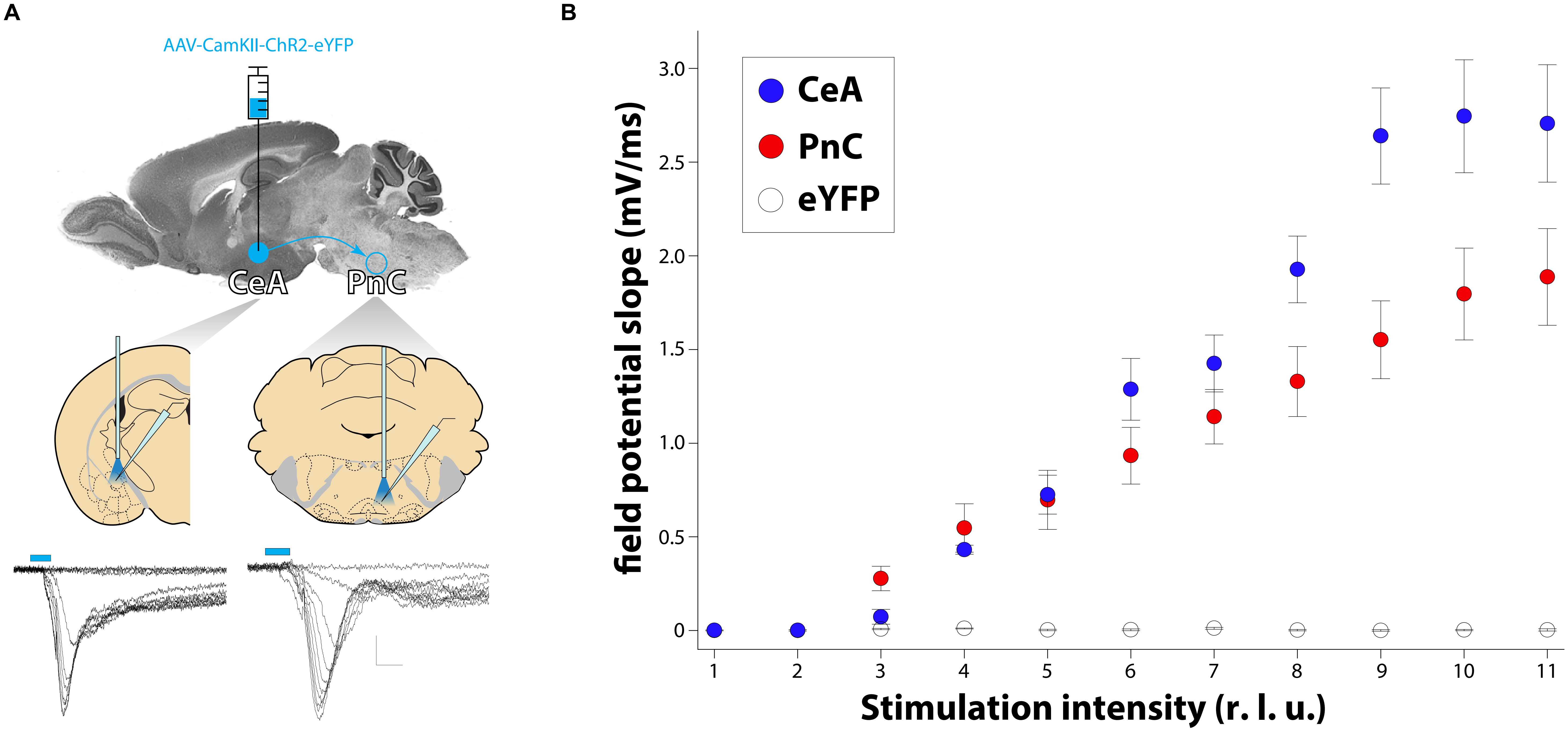

### Additional File 4

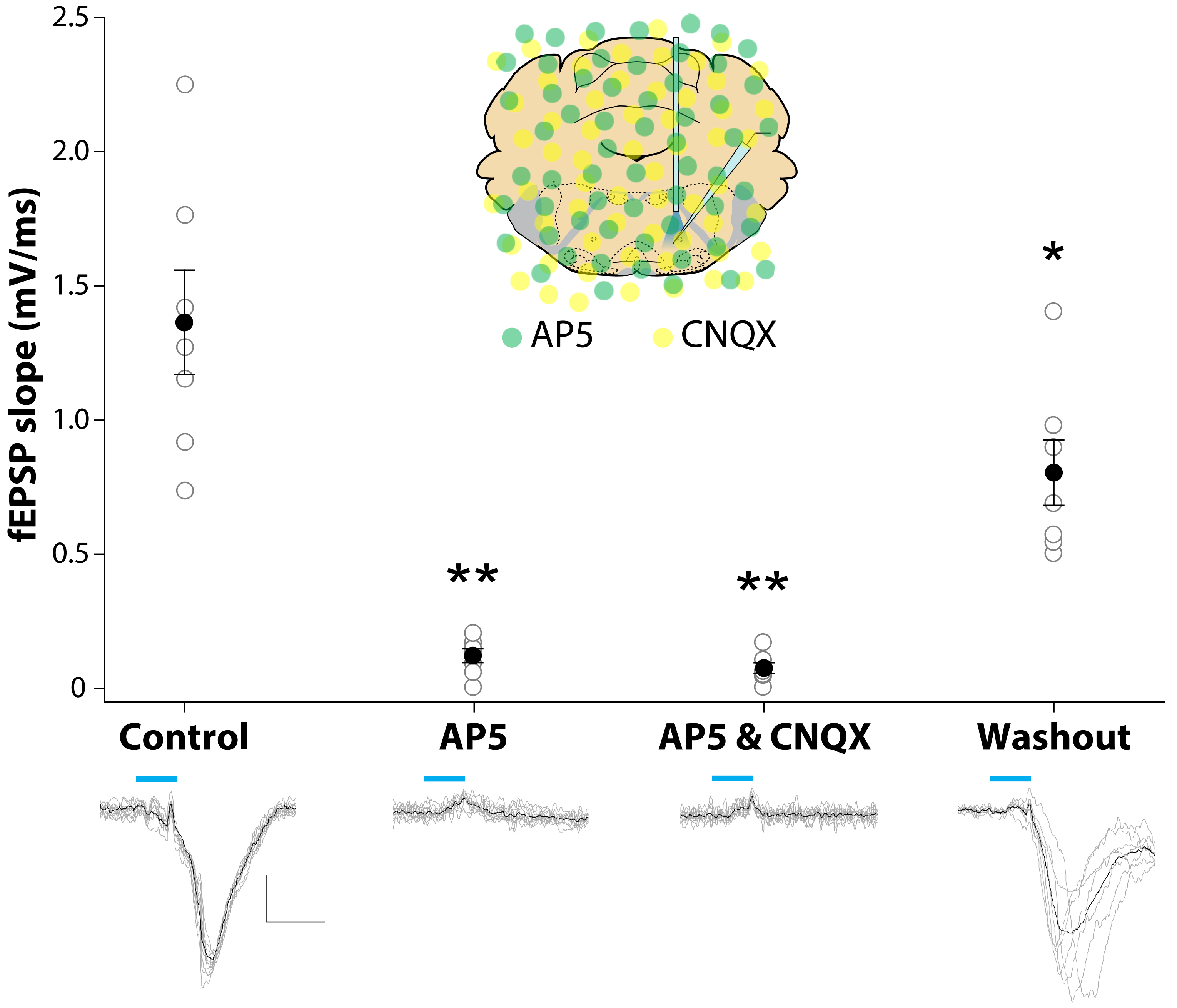

### Additional File 5

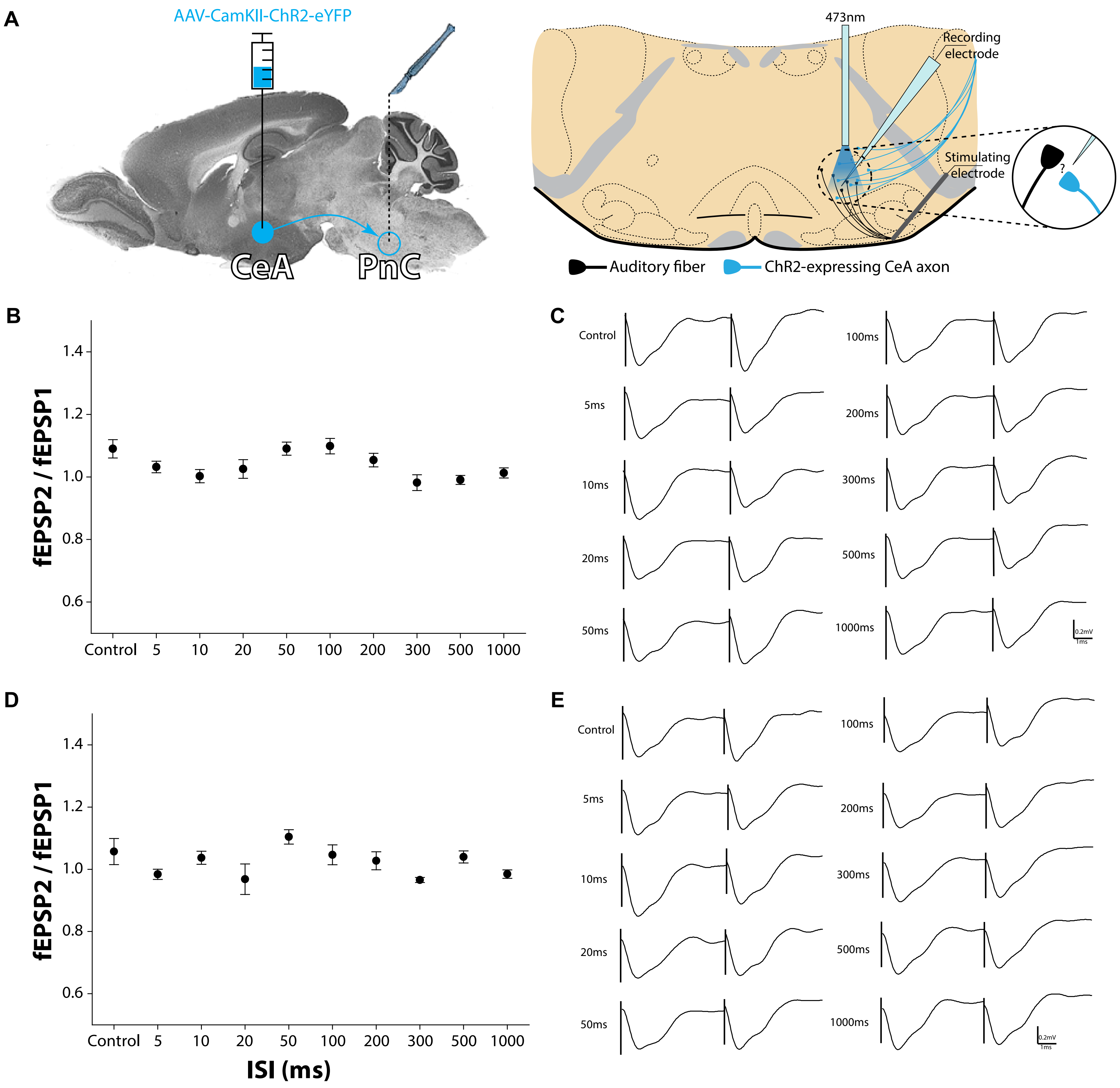

### Additional File 6

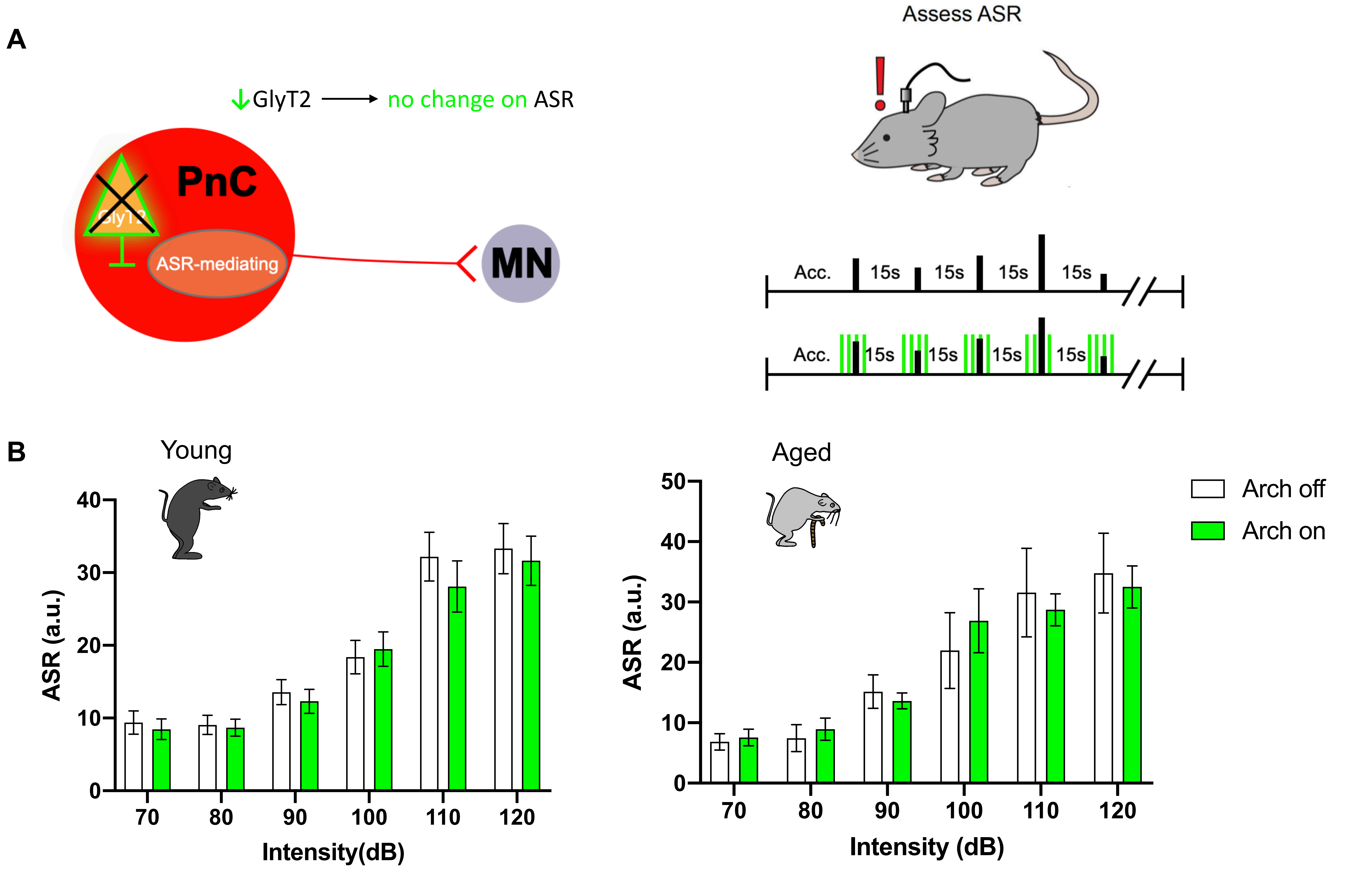

### Additional File 7

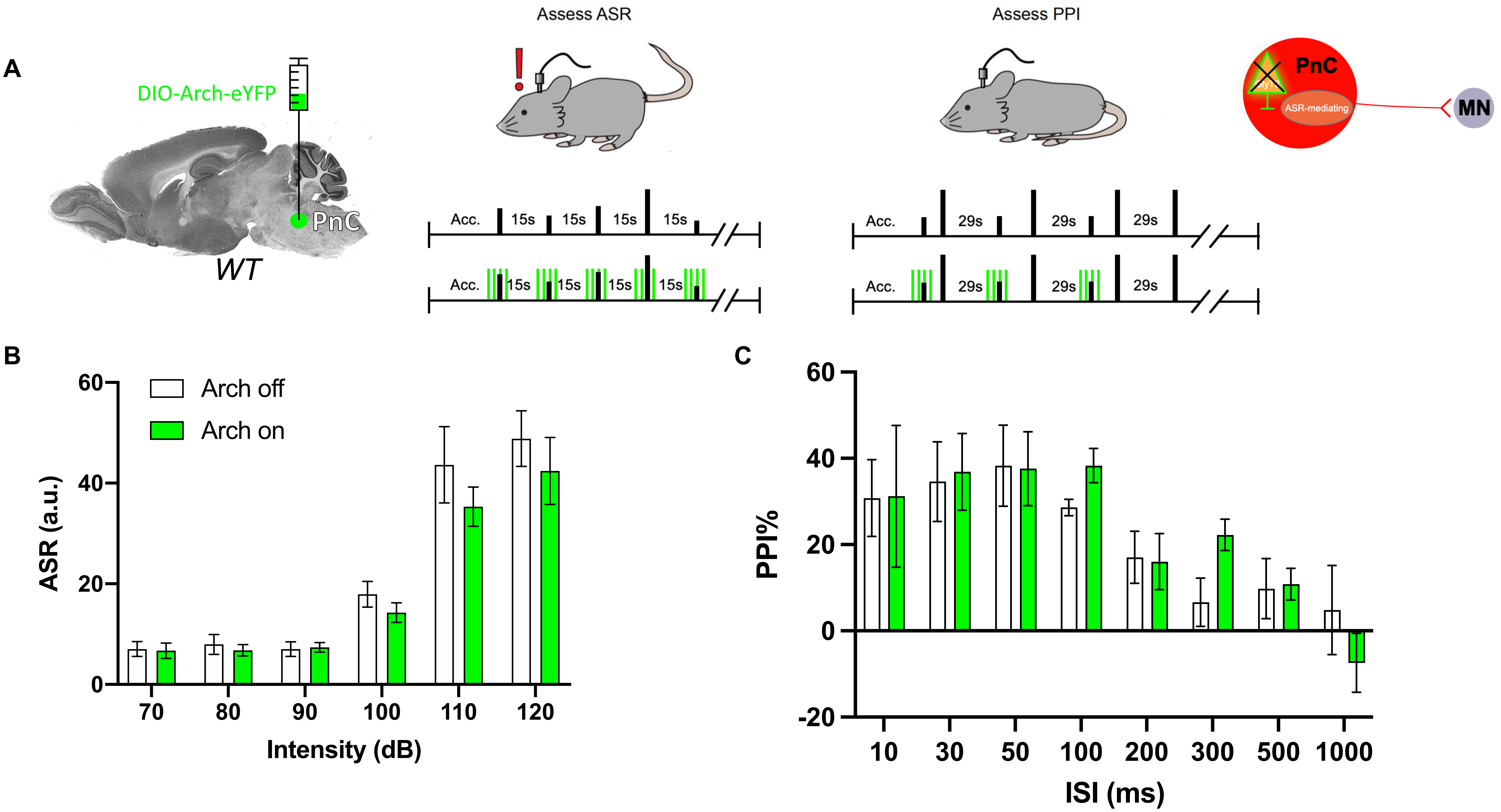
